# Reprogrammed CRISPR-Cas13b suppresses SARS-CoV-2 replication and circumvents its mutational escape through mismatch tolerance

**DOI:** 10.1101/2020.11.18.389312

**Authors:** Mohamed Fareh, Wei Zhao, Wenxin Hu, Joshua ML Casan, Amit Kumar, Jori Symons, Ilia Voskoboinik, Paul G Ekert, Rajeev Rudraraju, Sharon R Lewin, Joseph A Trapani

## Abstract

Mutation-driven evolution of SARS coronavirus-2 (SARS-CoV-2) highlights the need for innovative approaches that simultaneously suppress viral replication and circumvent viral escape routes from host immunity and antiviral therapeutics. Here, we employed genome-wide computational prediction and singlenucleotide resolution screening to reprogram CRISPR-Cas13b against SARS-CoV-2 genomic and subgenomic RNAs. Reprogrammed Cas13b effectors targeting accessible regions of Spike and Nucleocapsid transcripts achieved >98% silencing efficiency in virus free-models. Further, optimized and multiplexed gRNAs suppressed viral replication by up to 90% in mammalian cells infected with replication-competent SARS-CoV-2. Unexpectedly, the comprehensive mutagenesis of guide-target interaction demonstrated that single-nucleotide mismatches do not impair the capacity of a potent single gRNA to simultaneously suppress ancestral and mutated SARS-CoV-2 in infected mammalian cells, including the highly infectious and globally disseminated Spike D614G mutant. The specificity, efficiency and rapid deployment properties of reprogrammed Cas13b described here provide a molecular blueprint of antiviral therapeutics to simultaneously suppress a wide range of SARS-CoV-2 mutants, and is readily adaptable to other emerging pathogenic viruses.

## INTRODUCTION

Severe Acute Respiratory Syndrome Coronavirus 2 (SARS-CoV-2), the cause of coronavirus disease 2019 (COVID-19) has caused >52 million infections and over 1.3 million deaths worldwide as of November 2020^1^. A wide range of preventive and therapeutic strategies are the subject of intensive investigation and include vaccines^2^, engineered monoclonal antibodies^3,4^, and small molecule antiviral agents^5,6^. However, the capacity of viruses to evolve in response to host, environmental and therapeutic pressures presents challenges to these conventional approaches.

While SARS-CoV-2 possesses a moderate mutational rate due to the proofreading activity of its RNA-dependent RNA polymerase (RdRP), there are nonetheless reports of emerging SARS-CoV-2 strains that have evolved increased fitness and pathogenicity^7,8^. The SARS-CoV-2 D614G (Asp614Gly) variant, resulting from a single nucleotide substitution (A to G) in the receptor binding domain of the Spike glycoprotein, has demonstrated increased affinity for the ACE2 receptor and enhanced replication capacity in vitro, potentially contributing to the global spread of this variant^8–10^. Likewise, analysis of virus phylogenetics globally enabled by the GISAID^11^ and Nextstrain^12^ databases revealed recurrent hotspot mutations in several viral subunits that are considered likely to confer selective advantage and/or alter pathogenicity^13,14^. Also, there are concerns regarding uncontrolled spread and evolution of the virus in farm animals such as minks that could eventually lead to problematic mutations^15^. These findings emphasize the need for an innovative antiviral approach that can counteract dynamic changes in the SARS-CoV-2 genome and thwart viral escape.

The Clustered Regularly Interspaced Short Palindromic Repeats (CRISPR) Cas13 is a form of bacterial adaptive immunity that suppresses bacteriophage RNA^16–18^. Previous reports have demonstrated the potential of certain Cas13 orthologs to silence endogenous and viral RNAs in mammalian cells^19–21^, however, it remains to be established whether Cas13 is capable of silencing replication-competent SARS-CoV-2 in infected mammalian cells, and potentially, redesigned to suppress mutation-driven viral evolution.

## RESULTS

### *In-silico* prediction and design of a genome-wide SARS-CoV-2 gRNA library

We hypothesized that the SARS-CoV-2 RNA genome and its subgenomic transcripts can be silenced with CRISPR-Cas13 in mammalian cells. We opted to use pspCas13b^22^ due to its long (30-nt) spacer sequence that is anticipated to confer greater specificity than Cas13 orthologs with shorter target sequences. We used the first publicly available genomic sequence of SARS-CoV-2 as the reference genome^23^ to design pspCas13 gRNAs, and developed a bioinformatics pipeline for *in silico* design of 29,894 pspCas13 tiled gRNAs covering the entire genome, from 5’ to 3’ end (**Suppl. Figure 1 & Suppl. Table 1**). We further refined the list of gRNAs by excluding (i) spacer sequences with poly-T repeats (4 or more successive T’s) that prematurely terminate the Pol III-driven transcription of gRNAs (**Suppl. Table 2**), and (ii) spacers or target sequences with predicted thermodynamically stable RNA-RNA duplexes that should hinder gRNA loading into pspCas13b and target accessibility. We selected 839 spacer sequences that best satisfied the selection criteria (**Suppl. Table 3**). As all 839 30-nt spacer sequences fully base-pair with the reference SARS-CoV-2 RNA, all are predicted to achieve high targeting efficiency. This database represents a valuable investigative tool to interrogate SARS-CoV-2.

### PspCas13b suppresses the Spike transcript with high efficiency and specificity

In a proof-of concept assay, we used the bioinformatics pipeline described above to select gRNAs targeting the transcript of the Spike glycoprotein, which facilitates viral invasion of host cells following binding to the ACE2 surface receptor^3,24^ A codon-optimized Spike DNA template was cloned in frame with an upstream P2A selfcleavage peptide and enhanced green fluorescent protein (eGFP), enabling cotranscription and translation of Spike and eGFP, which are separated post-translationally by P2A proteolytic self-cleavage. In this reporter assay, pspCas13b-mediated cleavage of the Spike mRNA was predicted to lead to a loss of eGFP fluorescence (**Figure 1A**). We co-transfected 293 HEK cells with the Spike-eGFP reporter plasmid together with pspCas13 linked to a blue fluorescent protein (BFP) and gRNAs targeting either the Spike transcript (4 gRNAs) or non-targeting (NT) gRNA as a control. Fluorescence microscopy revealed that high silencing efficiency was achieved with all Spike-targeting gRNAs compared to the NT control (**Figure 1B & 1C**). We achieved similar silencing in VERO cells, a kidney epithelial cell line derived from an African green monkey, commonly used in SARS-CoV-2 research due to its high susceptibility to infection^25^ (**Figure 1B & 1C**). gRNA2 achieved the highest silencing efficiency amongst the 4 gRNAs we tested, reaching >99% and >90% reduction in spike transcript levels in 293 HEK and VERO cells, respectively. To demonstrate that the observed silencing was dependent on the cellular expression of gRNA, we transfected 293 HEK cells with increasing amounts of either NT or Spiketargeting gRNA2 plasmids ranging from 0 to 104 pM (0, 0.16, 0.8, 4, and 20ng plasmid in 100 μL volume). While NT gRNA exhibited no effect on eGFP expression, the Spiketargeting gRNA2 showed dose-dependent silencing of the Spike transcript with 50% silencing efficiency (IC_50_) achieved with 5.16 pM of plasmid gRNA (equivalent to 994pg in 100 μL of media), demonstrating that gRNA availability in the cell is key for efficient degradation of viral RNA (**Figure 1D & 1E**). This very low IC_50_ value also highlights the need for intracellular delivery of just a few copies of gRNA template plasmid to achieve effective silencing, thanks to intracellular gRNA amplification by the transcription machinery.

**Figure 1.**
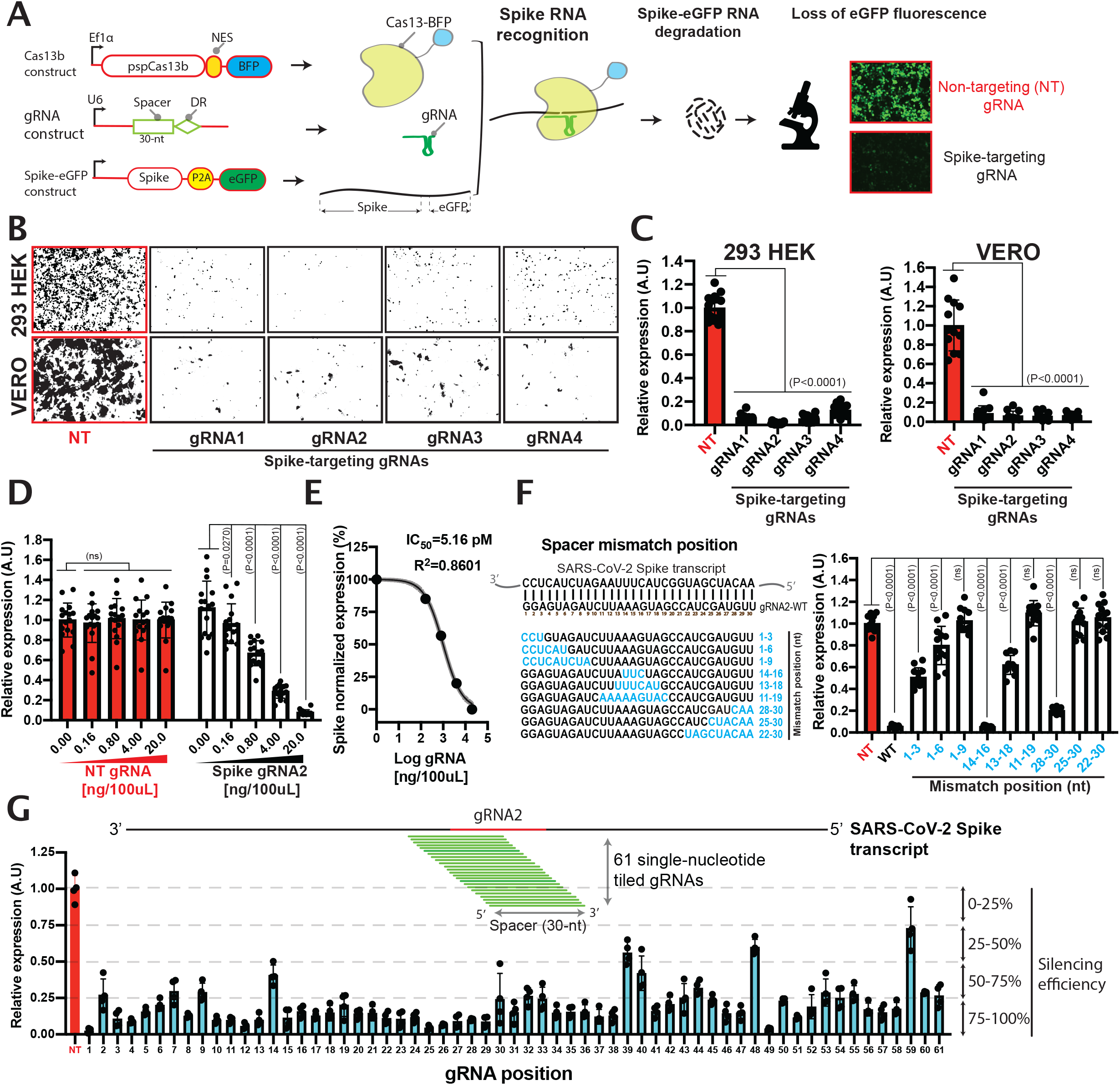
Reprogrammed PspCas13b silences the Spike transcript with high efficiency and specificity. (**A**) Schematic of pspCas13b reporter assay used to track the recognition and degradation of SARS-CoV-2 Spike RNA. (**B**) Representative fluorescence microscopy images show the silencing of the Spike transcript with 4 targeting gRNAs in 293 HEK (upper panel) and VERO cells (lower panel). NT is a non-targeting control gRNA; Images are processed for quantification using ImageJ. (**C**) Quantification of silencing efficiency with various gRNAs in 293 HEK and VERO cells (4 representative field of views were imaged per condition; N=3). (**D**) gRNA dosedependent silencing of the Spike transcript with either NT (red) or spike targeting gRNA2 (black) in 96-well containing 100 μL of media; N=4. (**E**) Spike gRNA2 IC_50_ value to silence 50% of the Spike transcript. (**F**) Mutagenesis analysis of spacer-target interaction. The nucleotides in blue highlight residues in the spacer sequence that were mismatched with the targeted sequence. N=4. Results analysed with one-way Anova test (95% confidence interval). (**G**) Tiling of 61 gRNAs with single-nucleotide increment reveals that RNA landscape influences pspCas13b silencing efficiency. Data are normalized mean fluorescence per field of view, and errors are SD. N=1 with four representative field of views. N is the number of biological replicates

Like all RNA-guided nucleases, pspCas13b specificity is conferred by basepairing between the target and gRNA spacer sequences^26–28^. Evaluating the tolerance for mismatches in this interaction is therefore critical to determining both the potential off-target activity of pspCas13b (ie, silencing cellular transcripts), and more importantly, the loss of activity against variant sequences generated by the RdRP during viral replication^29,30^. Given that pspCas13b specificity is still poorly understood, we opted to determine the degree of variation in base-pairing between target RNA and the spacer of gRNA2 that could be tolerated and still lead to RNA degradation. We designed and cloned 9 constructs that encoded successive 3-, 6-, or 9-nt substitutions across the whole gRNA2 spacer sequence (**Figure 1F**). The nucleotide substitutions created mismatches (MSM) that perturbed the thermodynamic stability of the RNA-RNA duplex. While a 3-nt mismatches placed internally (position 14-16) or at the 3’end (position 28-30) had minimal impact, 3-nt mismatches at the 5’end reduced the silencing efficacy by ~50%. By comparison, 6-nt mismatches introduced at various locations markedly and consistently reduced silencing, and 9-nt mismatches completely abolished the degradation of the Spike transcript (**Figure 1F**). Taken together, this experiment indicated that gRNA2 requires >21-nt base-pairing with its target to trigger the necessary CRISPR-Cas conformational change and nuclease activation necessary for target degradation^26–28^. Conversely, the ability of pspCas13b to tolerate up to 3-nt mismatches underlines its potential to remain effective against the majority of variants with single-nucleotide polymorphisms in the target sequence, conferring protection against potential viral escape mutants such as the D614G mutation in the SARS-CoV-2 Spike protein^8^, and mutations that compromise the efficacy of therapeutic antibodies against SARS-CoV2^7^.

Next, we questioned whether the RNA target sequence and structure may influence pspCas13b silencing efficiency. To comprehensively evaluate this possibility, we generated 61 single-nucleotide-tiled gRNAs covering the 5’ and 3’ overhangs of the gRNA2 target sequence (**Figure 1G**). This approach generated single-nucleotide resolution of the RNA landscape that influences pspCas13b silencing. Overall, 95% of gRNAs achieved >50% degradation efficiency, and 70% of gRNAs achieved >75% degradation of the Spike transcript. Notably, the least effective gRNAs were spatially clustered, suggesting that the RNA secondary structure and/or the presence of endogenous RNA binding proteins might limit spatial RNA accessibility. The tiling approach employed results in continuous changes in spacer flanking sequences (PFS). Accordingly, our findings revealed that unlike other CRISPR subtypes, pspCas13b activity appears to be independent of defined PFS or PAM-like (protospacer adjacent motif) motifs that constrain the targeting spectrum of other CRISPR effectors^31^. These data highlight the high likelihood of efficient silencing with various pspCas13b gRNAs and emphasize the flexibility of this RNA silencing tool for SARS-CoV-2 suppression.

### Silencing SARS-CoV-2 nucleocapsid transcript

We next examined whether other SARS-CoV-2 structural components essential for viral assembly can be silenced by pspCas13b. We chose to target the RNA encoding the nucleocapsid protein (NCP) that is critical for viral particle assembly by packaging the RNA genome within the viral envelope^32,33^. Based on the reference genome sequence^23^ we designed and cloned the original sequence of the NCP into a reporter vector in frame with a monomeric red fluorescent protein (mcherry) (**Figure 2A**). We co-transfected 293 HEK and VERO cells with the NCP-mcherry reporter plasmid together with pspCas13-BFP and a NT gRNA or three different gRNAs targeting various regions of the NCP transcript.

**Figure 2.**
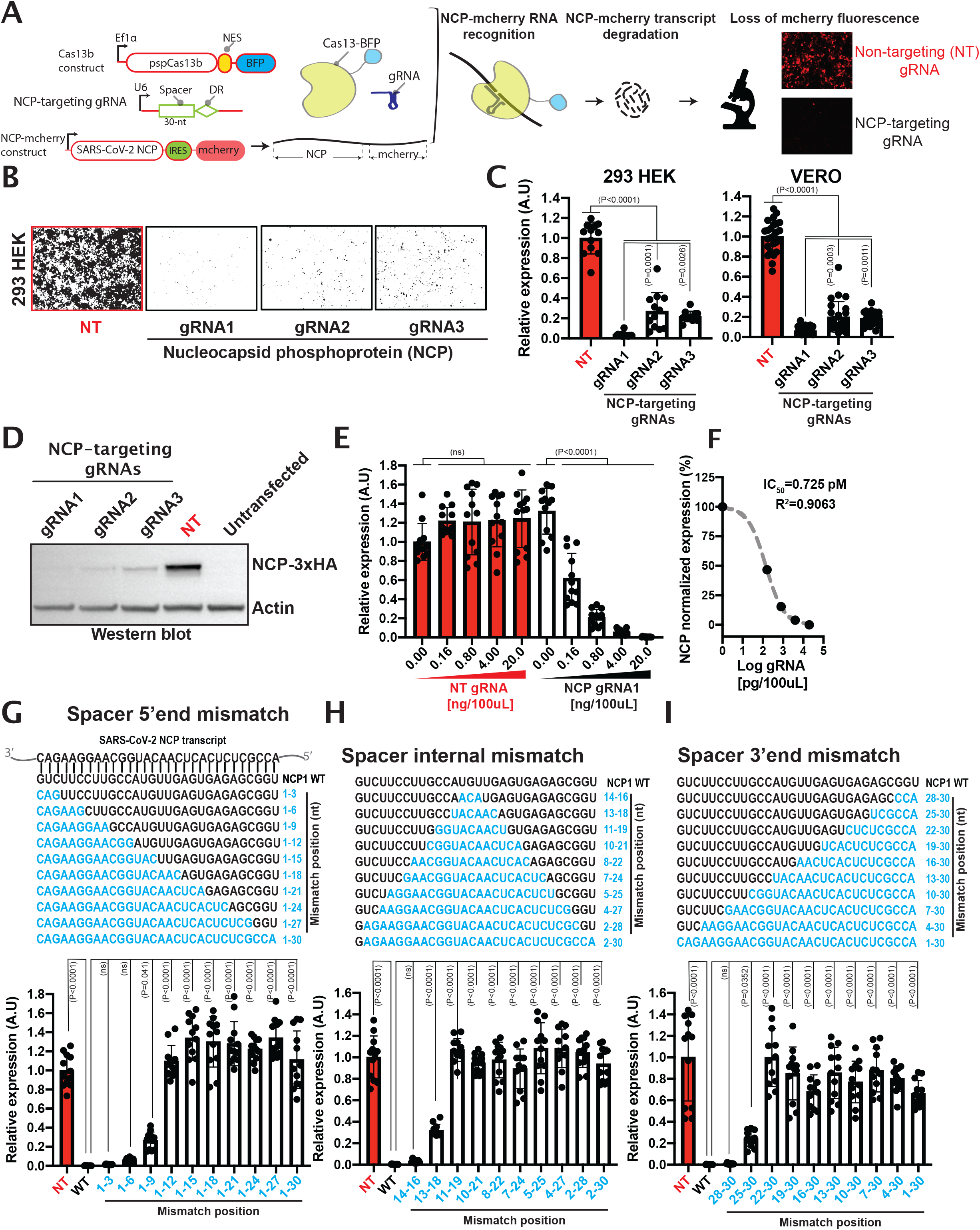
PspCas13b suppresses SARS-CoV-2 nucleoprotein (NCP) transcript with high efficiency and specificity. (**A**) Schematic of pspCas13b reporter assay to monitor NCP silencing efficiency of various gRNAs. (**B**) Representative fluorescence microscopy images show the silencing efficiency of the NCP transcript with 3 targeting gRNAs in 293 HEK. NT is a non-targeting control gRNA. (**C**) The histograms quantify the silencing efficiency of various gRNAs in 293 HEK (N=3) and VERO cells (N=6). (**D**) Western blot analysis of NCP silencing efficiency in 293 HEK obtained with various gRNAs (N=3). (**E**) gRNA dose-dependent silencing of the NCP transcript with either NT (red) or NCP targeting gRNA1 (black) in 96-well containing 100 μL of media; (N=3). (**F**) NCP gRNA1 IC_50_ value (pg) to silence 50% of the Nucleocapsid transcript. (**G-I**) Comprehensive analysis of spacer-target interaction, examining specificity and mismatch tolerance of papsCas13b gRNA at various positions of the spacer. The nucleotides in blue highlight residues in the spacer sequence in which mismatches with the targeted sequence were introduced. Microscopy data are normalized mean fluorescence and errors are SD. 4 representative field of views per condition were imaged in N=3. N is the number of biological replicates. Results analysed with one-way Anova test (95% confidence interval); N is the number of biological replicates

Based on mcherry fluorescence intensity, we achieved high silencing efficiency in both cell lines, with all three NCP-targeting gRNAs (**Figure 2B & 2C**). Of these, gRNA1 exhibited the highest silencing efficiency, achieving >99% and >90% reduction in fluorescence in 293 HEK and VERO cells, respectively. In line with the microscopy data, western blot confirmed that the three NCP-targeting gRNAs efficiently depleted NCP protein, with gRNA1 again the most effective (**Figure 2D**). As with Spike-targeting gRNAs, titration of the NCP-targeting gRNA1 plasmid (0, 0.83, 4.15, and 20.75 pM, equivalent to 0, 0.16, 0.8, 4, and 20ng per 100 μL of media) into 293 HEK cells demonstrated dose-dependent silencing of the target, whereas NT gRNA had no effect (**Figure 2E**). The dose-dependent effect of NCP-targeting gRNA1 revealed 50% inhibition of the target (IC_50_) with 0.725 pM of plasmid gRNA (140 pg in 100 μL of media in 96-well format; R^2^=0.906) (**Figure 2F**), whereas the IC_50_ of Spike-targeting gRNA2 was 5.16 pM (R^2^=0.86) (**Figure 1E**). The variation in silencing efficiency (7-fold) observed with these two potent spacer sequences again highlighted that affinity and target accessibility need to be considered when selecting the optimal gRNA for maximal target silencing.

Next, we examined the extent to which mismatches in the spacer-target RNA-RNA hybrid of the highly efficient NCP-targeting gRNA1 would compromise its silencing potency. We designed 29 additional gRNA constructs that incorporated mismatches at the 5’ end, 3’ end, or at internal positions (blue residues, **Figure 2G-I**). Mismatch length varied from 3 to 30 nucleotides to cover the entire spacer sequence. At the 5’end, mismatches at positions 1-3 or 1-6 were well-tolerated and caused minor reductions in silencing efficiency in 293 HEK cells transfected with 20ng of gRNAs, while altering positions 1-9 reduced silencing by ~30%. However, by titrating the quantity of gRNA delivered, we revealed noticeable loss of silencing efficiency when 1-3, 1-6 or 1-9 mismatches were introduced (**suppl. Figure 2**). Mismatches longer than 9-nt at the 5’end completely abrogated silencing. Similarly, 3-nt mismatches placed internally (positions 14-16) or at the 3’ end were well tolerated, while 6-nt mismatches reduced silencing by ~30%. Introducing >6-nt mismatches at both internal and 3’ end positions caused complete loss of silencing (**Figure 2H & 2I**).

### pspCas13b silencing tolerates a single-nucleotide mismatch with the target

Viral evolution takes place predominantly through single-nucleotide indels brought about by error-prone polymerases. We investigated whether pspCas13b can tolerate a single-nucleotide mismatch between the spacer and the SARS-CoV-2 RNA target, potentially enabling our silencing technology to remain effective against spontaneous mutations. We re-designed the gRNAs targeting either the Spike (gRNA2) or the nucleocapsid (gRNA1) transcripts to harbour a single-nucleotide mismatch at spacer positions 1, 5, 10, 15, 20, 25, and 30 and compared their silencing efficiency to their fully-matched wildtype counterparts. Overall, we found that a single-nucleotide mismatch at various locations of the spacer was well tolerated, with no appreciable impact on target silencing (**Figure 3A & 3B**). Among all the constructs we tested, only a single nucleotide mismatch in the first nucleotide of the spacer sequence of gRNA1 targeting Nucleocapsid exhibited a moderate loss of silencing (**Figure 3B**). In line with the fluorescence data, western blot analysis of Nucleocapsid protein expression confirmed that single-nucleotide mismatch at position 5, 10, 15, 20, 25, and 30 are well tolerated and retain full silencing efficiency, while a mismatch at position 1 led to a partial loss of silencing (**Figure 3C**). These data highlight the potential of a single pspCas13b gRNA to silence SARS-CoV-2 variants and overcome mutation-driven viral escape.

**Figure 3.**
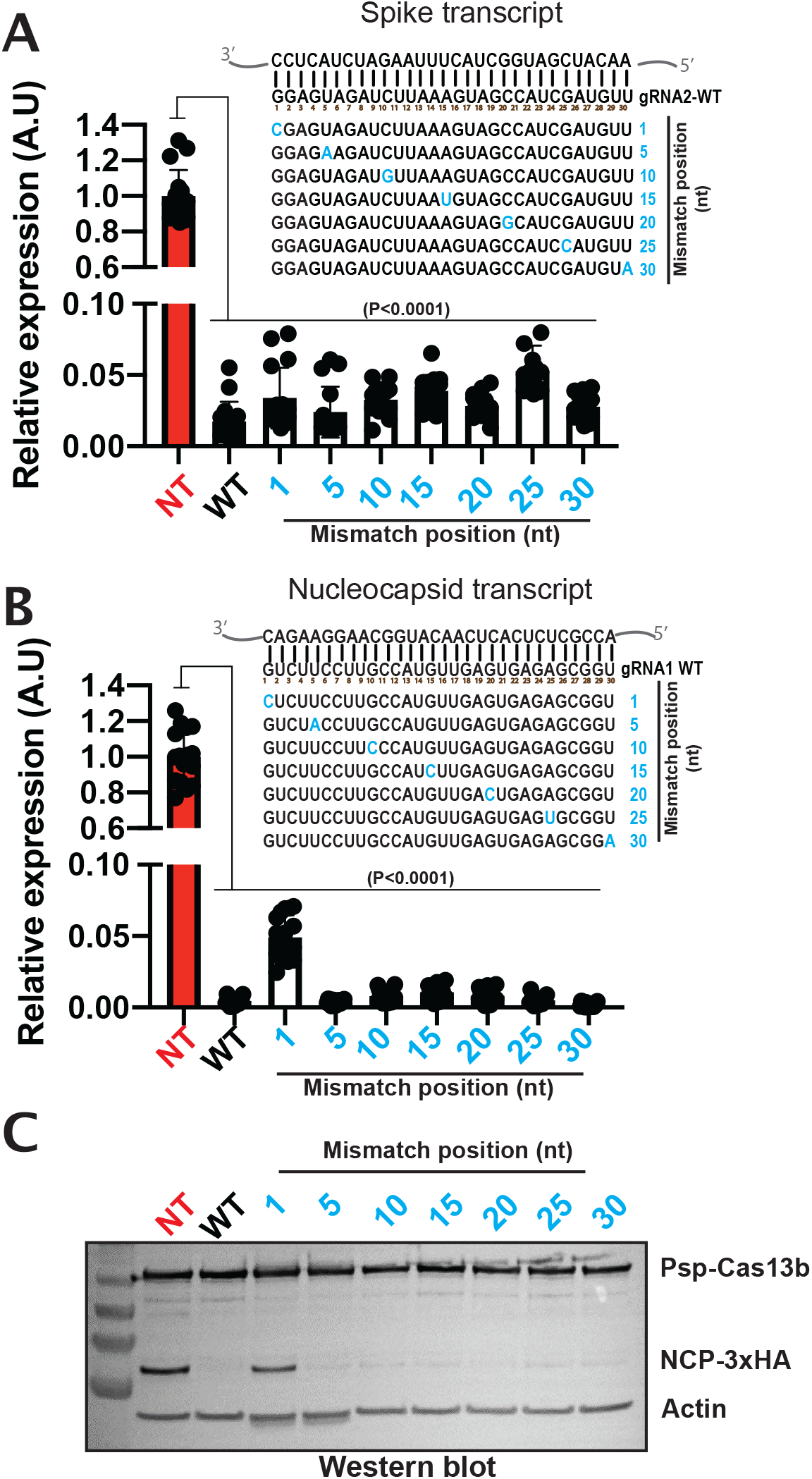
PspCas13b silencing tolerates single-nucleotide mismatch with SARS-CoV-2 targets. Fluorescence-based reporter assays to assess the silencing efficiency of the Spike (**A**) or Nucleocapsid (**B**) transcripts with gRNAs harbouring singlenucleotide mismatch at various location of the spacer-target interface 48h posttransfection. 4 representative field of views per condition were imaged; N=4. Fluorescence data are normalized means and errors are SD. Results are analysed by one-way ANOVA test (95% confidence interval). (**C**) Representative Western blot analysis to examine the expression level of Nucleocapsid protein (3xHA tagged) in 293 HEK cells expressing NCP-1 gRNA with single-nucleotide mismatch at various positions of the spacer-target interface 24h post-transfection; N=3. N is the number of biological replicates.

### PspCas13b reduces live SARS-CoV-2 virus replication

A recent study attempted to target a SARS-CoV-2 RNA construct with a Cas13d ortholog but the silencing of live replication-competent SARS-CoV-2 was not examined^21^. Rather, a genetically engineered H1N1 influenza strain expressing a fluorescent reporter was used. To determine whether our reprogrammed pspCas13b could inhibit SARS-CoV-2 RNA replication, we transfected VERO cells with pspCas13b-BFP and either NT gRNA or the gRNA1 that most efficiency reduced Nucleocapsid RNA levels in virus-free models (**Figure 2**). We optimised transfection conditions of VERO cells to achieve 25-40% transfection efficiency as measured by flow cytometry analysis (**Suppl. Figure 3**). 48h post-transfection, we infected the cells with replication-competent SARS-CoV-2 at a Multiplicity Of Infection (MOI) of 0.01 or 0.1, and assayed the culture medium at 1hr, 24hrs, and 48hrs time-points using real-time PCR (RT-PCR) for viral RNA^25^. We also determined the titre of infectious SARS-CoV-2 by exposing fresh VERO cells to the supernatant and quantifying the cytopathogenic effect (CPE) by light microscopy and manual counting to determine the tissue culture infectious dose that inhibits 50% of virus growth (TCID50) (**Figure 4A**). In cells transfected with NT gRNA and infected at 0.1 MOI, viral RNA in the supernatant increased ~15-fold at 24h compared to baseline and this level was maintained at 48hrs. By contrast, in the presence of NCP-targeting gRNA1 there was a marked reduction in viral RNA detected (**Figure 4B**). RT-PCR analysis showed that gRNA1 reduced the viral load by ~90% 24h post-infection compared to the NT gRNA. However, NCP-targeting gRNA1 did not completely abrogate viral replication in this biological replicate, as the viral titre still increased 3-fold between 24 and 48hrs (**Figure 4B**). This partial inhibition of SARS-CoV-2 replication by pspCas13b is likely explained by a combination of a high viral load (0.1 MOI) that saturates intracellular pspCas13b nucleoproteins, and the limited transfection efficiency in VERO cells (25-40%) (**Suppl. Figure 3**). Subsequently, viral replication may take place in untransfected cells or cells expressing low levels of pspCas13b and gRNA. Increasing the efficiency of delivery of pspCas13b and its gRNA should result in even greater viral suppression.

**Figure 4.**
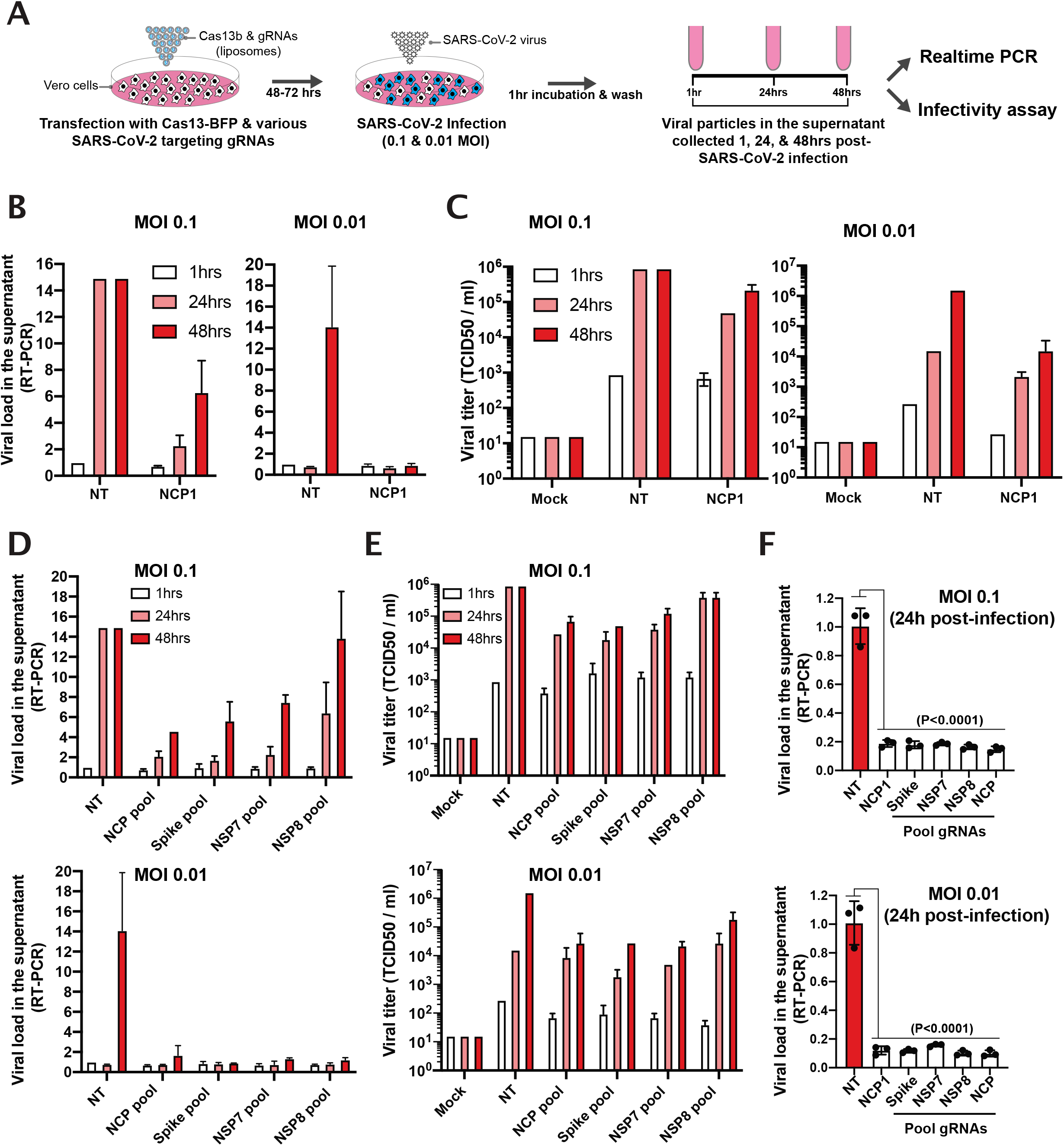
Silencing of SARS-CoV-2 virus replication in VERO cells. (**A**) Schematic of infection assay to assess pspCas13b-mediated suppression of SARS-CoV-2 replication in VERO cells. VERO cells were transfected with liposomes containing pspCas13b-BFP and various gRNA constructs. 48-72h post-transfection, cells were infected with SARS-CoV-2 for 1 hour and the kinetics of viral replication was assessed in supernatants collected at 1, 24, and 48h post-infection for production of viral RNA using RT-PCR and infectious virus by quantification of the TCID50 using limiting dilution infection of VERO cells and detection of CPE. (**B**) Representative RT-PCR and (**C**) infectivity assay to evaluate the kinetic of viral replication in VERO cells expressing either NT or NCP-targeting gRNA1 48h post-transfection of pspCas13b and gRNAs transfection, N=2. (**D**) Representative RT-PCR and (**E**) infectivity assays to monitor the kinetics of viral replication in VERO cells expressing either NT or various pools of gRNA targeting NCP, Spike, NSP7, and NSP8. VERO cells were infected with a replication-competent SARS-CoV-2 48h post-transfection of pspCas13b and gRNAs; N=2. (**F**) RT-PCR analysis of the viral load in the supernatant of VERO cells expressing either NT, NCP-1, or various pools of gRNA targeting NCP, Spike, NSP7, and NSP8. VERO cells were infected with a replication-competent SARS-CoV-2 48h post-transfection of pspCas13b and gRNAs; N=3; Data are normalized means and errors are SD; Results analysed with one-way Anova test; N is the number of biological replicates (95% confidence interval).

In fact, when we infected VERO cells with one order of magnitude lower viral titre (0.01 MOI), viral RNA in the supernatant of cells expressing NT gRNA increased by ~14-fold 48hrs post-infection. By contrast, viral RNAs in the culture medium of cells expressing NCP-1 gRNA remained at the basal level (**Figure 4B**), demonstrating that pspCas13b and NCP-targeting gRNA1 efficiently suppressed viral replication in infected mammalian cells. In line with RT-PCR, the infectivity assay confirmed that the NCP-targeting gRNA1 greatly reduced the release of infectious virus (measured as TCID50 / mL) in the supernatant (**Figure 4C**).

The emergence of new viral strains of SARS-CoV-2, potentially with enhanced fitness, poses a formidable challenge to any single antiviral or antibody therapy^7^. Likewise, targeting SARS-CoV-2 with a single gRNA may have limited efficacy due to viral adaptation and/or poor accessibility to viral RNA *in vivo* due to intrinsic RNA folding and/or interference from RNA-binding proteins. We postulated that simultaneously targeting several regions of SARS-CoV-2 viral RNAs with multiplexed gRNAs would mitigate these effects, and reduce the likelihood of viral adaptation. We transfected VERO cells with four pools of gRNAs each containing four different guides targeting either SARS-CoV-2 structural proteins Spike and NCP, or non-structural protein NSP7 and NSP8 that act as co-factors critical for the RdRP^6,29,30^. We again infected the VERO cells with SARS-CoV-2, and quantified viral RNA and infectious virus in the culture supernatant. We found that all four gRNAs pools markedly reduced viral RNA (**Figure 4D**), and infectious virus (**Figure 4E,**) in the supernatant at both 0.1 and 0.01 MOI.

Likewise, when we challenged VERO cells with 0.1 and 0.01 MOI of SARS-CoV-2 virus 72h post-transfection of pspCas13b and various gRNAs, we achieved ~80% and ~90% suppression of viral loads with all tested gRNAs, respectively (**Figure 4F & Suppl. Figure 4**). These results indicate that SARS-CoV-2 viral suppression from a single pspCas13b transfection can persist beyond 5 days post-transfection, and 3 days post-infection.

To the best of our knowledge, this is the first study that definitively demonstrates the effective and specific targeting of replication-competent SARS-CoV-2 in infected mammalian cells, using CRISPR-Cas13.

### pspCas13b suppresses mutation-driven SARS-CoV-2 evolution

A single nucleotide substitution in the receptor binding domain (RBD) of the Spike protein led to the global emergence of SARS-CoV-2 D614G variant with increased ACE-2 affinity and infective potential^8,9^. Based on the single-nucleotide mismatch data described above (**Figure 3A & 3B**), we hypothesized that gRNAs designed against ancestral Spike D614 genomic sequence should remain effective against the D614G mutant RNAs despite a single-nucleotide mismatch at the spacer-target interface. To test this, we cloned part of the D614G Spike coding sequence (1221 bp; D614G SARS-CoV-2 genomic region between 22,760—23,980) into the mcherry reporter system, and assessed the silencing efficiency of 6 tiled gRNAs targeting this frequently mutated hotspot. The spacer sequences of all 6 tiled gRNAs tested are designed to fully match the ancestral Spike sequence (D614) and harbour one nucleotide mismatch with the Spike transcript of D614G mutant at spacer positions 5, 10, 15, 20, 25, and 30 (**Figure 5A**). The data showed that all 6 gRNAs significantly degraded the D614G mutant transcript (P<0.0001). Interestingly, gRNAs with a single-nucleotide mismatch at spacer position 15, 20, 25, or 30 were highly effective and exhibited >98% silencing efficiency against the D614G mutant transcript. By contrast, gRNAs with a mismatch at position 5 or 10 showed a reduced silencing efficiency estimated at approximatively 85-88% removal of the D614G mutant transcript (**Figure 5A**). These data suggest that pspCas13 and the D614G targeting gRNAs with a single-nucleotide mismatch at positions 15, 20, 25, and 30 are likely to remain effective against both ancestral and D614G mutant in infected cells. Thus, the mismatch tolerance molecular mechanism described here is expected to confer pspCas13d resilience against spontaneous single-nucleotide mutations that drive viral escape in SARS-CoV-2 and other pathogenic viruses.

**Figure 5.**
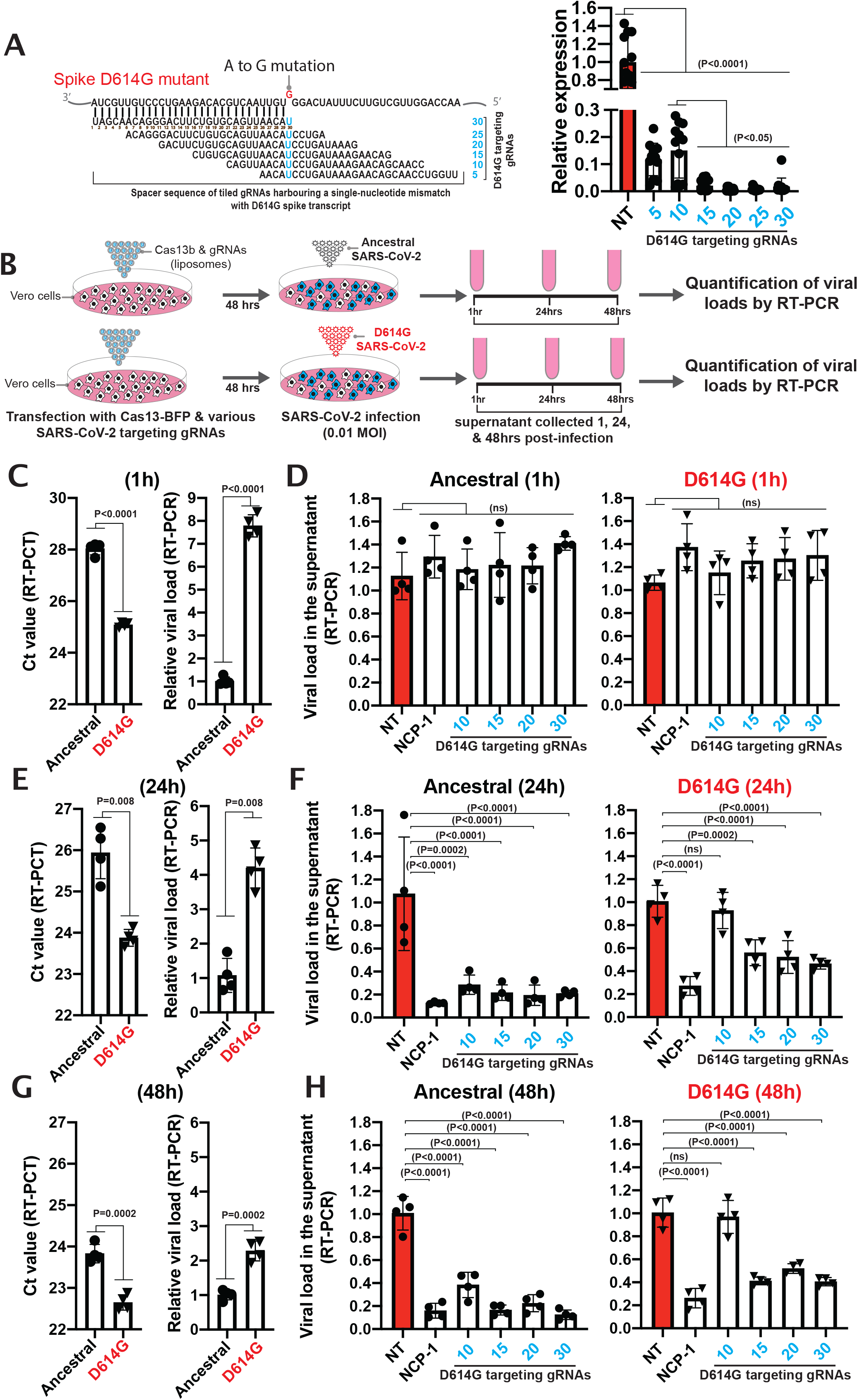
Single gRNAs targeting the D614 genomic mutation hotspot can silence both ancestral and D614G mutant due to single-nucleotide mismatch tolerance. **(A)** Fluorescence-based reporter assay to assess the silencing efficiency of the D614G Spike transcript with 6 tiled gRNAs harbouring a single-nucleotide mismatch with the target at various spacer positions, N=3; Fluorescence data are normalized means and errors are SD; Results are analysed by one-way ANOVA test (95% confidence interval). (**B**) Schematic of infection assay to assess pspCas13b-mediated suppression of both ancestral and D614G mutant in VERO cells. VERO cells were transfected with pspCas13b-BFP and various gRNA constructs. 48h post-transfection, cells were infected with either ancestral or D614G SARS-CoV-2 for 1 hour, and the kinetics of viral replication was assessed in supernatant collected at 1, 24, and 48h post-infection via quantification of viral RNA using RT-PCR. (**C,E, & G**) RT-PCR assays to evaluate relative viral loads (ancestral versus D614G) in control VERO cells expressing non-targeting (NT) gRNA at timepoints 1hr to estimate the initial viral input (**C**), 24h (**E**), and 48h (**E**) post-infection, N=4; Data are normalized means and errors are SD; Results are analysed by unpaired Student’s t-test (95% confidence interval). (**D,F, & H**) RT-PCR assays to evaluate the suppression of ancestral and D614G SARS-CoV-2 strains in VERO cells expressing either non-targeting (NT), NCP-1, or tiled gRNAs targeting the D614G mutation hotspot in the Spike genomic and subgenomic RNA 1hr (**D**), 24h (**F**), and 48h (**H**) post-transfection, N=4; Data are normalized means and errors are SD; Results are analysed by one-way ANOVA test (95% confidence interval).

**Figure 6.**
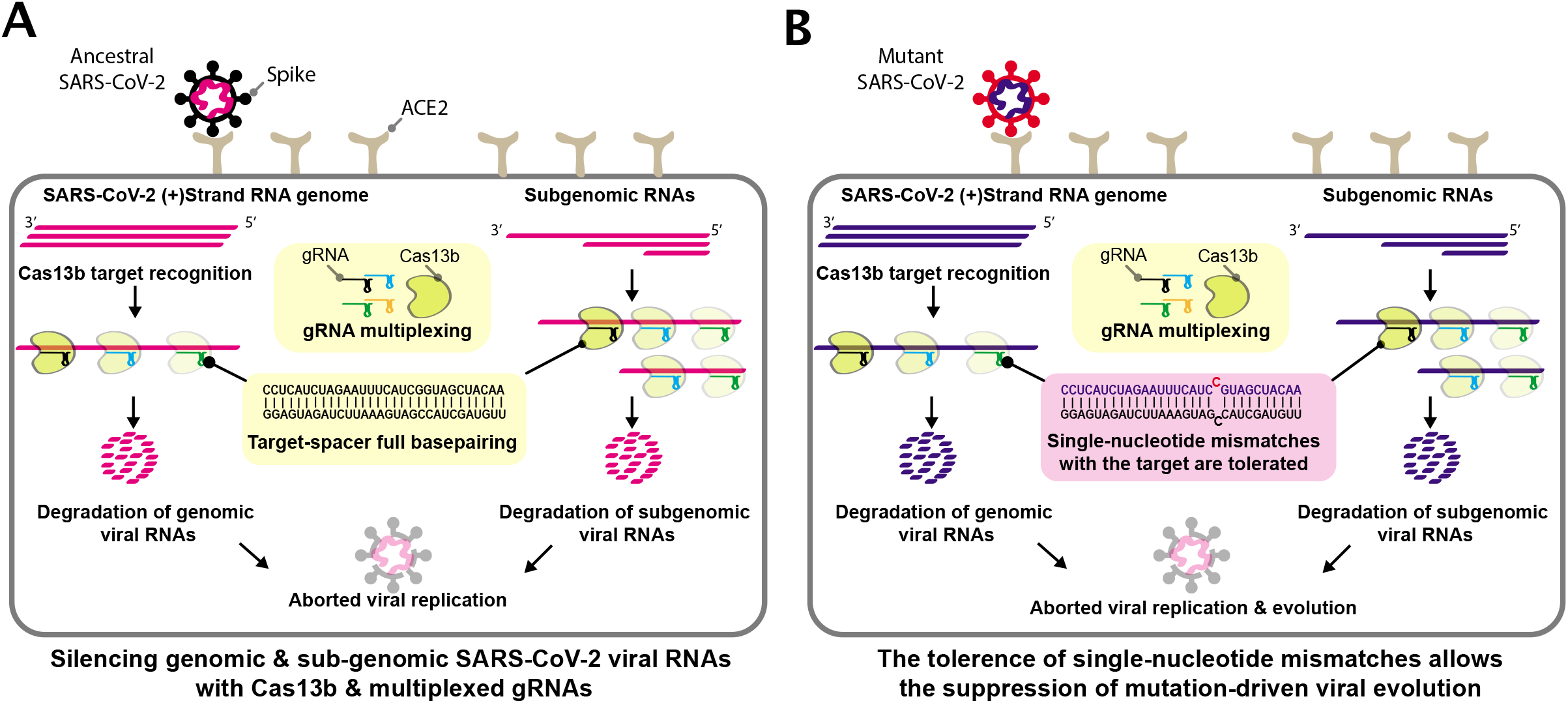
Schematic depiction of pspCas13b-mediated suppression of SARS-CoV-2 replication cycle and mutation-driven evolution through gRNA multiplexing and single-nucleotide mismatch tolerance. (**A**) gRNA multiplexing enables simultaneous targeting of several genomic and sub-genomic RNA locations, which limits the probability of poor silencing due to target inaccessibility (e.g. RNA folding, interaction with RNA-binding proteins *in vivo).* The spacer of pspCas13b mediates sequence-specific target recognition through complete base-pairing with the targeted sequence, followed by nuclease activation and target degradation. The silencing of viral RNAs alters the integrity of the viral genome, suppresses sub-genomic RNA-dependent viral protein translation, and the assembly of replication-competent viral particles within infected mammalian cells. This viral RNA targeting approach aborts viral replication cycles and inhibits the infectivity of SARS-CoV-2 virus. (**B**) The error-prone RdRP creates viral polymorphisms through the incorporation of single-nucleotide indels, and together with host (immune system) and environmental (antiviral therapies) positive pressures, can result in the emergence of new strains with higher infectivity and pathogenicity. Unlike other antiviral therapeutics (e.g. monoclonal antibodies^7^), pspCas13b target affinity remains unaffected when a single-nucleotide indel occurs at the target site, enabling a single gRNA to be effective against wild type and mutant strains. Thus, the pspCas13b approach can suppress viral evolution.

To test the mismatch tolerance of our Cas13b system against various replication-competent SARS-CoV-2 viral strains, we transfected VERO cells with NT gRNA, NCP-1 targeting gRNA (positive control), or 4 tiled gRNAs (10, 15, 20, 30) fully matching the ancestral Spike sequence D614 (see the schematic illustration in **Figure 5A & 5B**). 48 hours post-infection, VERO cells expressing pspCas13-BFP and various gRNAs were infected with either the ancestral or D614G mutant SARS-CoV-2, and the viral loads in supernatants were quantified by RT-PCR 1h (initial viral input), 24h and 48h post-infection (**Figure 5B**). The comparison of viral loads in the supernatant of cells infected with either the ancestral or D614G in the control groups (NT) showed 7.7, 4.2, and 2.3-fold higher viral loads in the D614G samples at timepoints 1h, 24h, and 48h, respectively (**Figure 5C, 5E & 5G**). These data revealed that the initial D614G viral load used here was more than 7 times higher than the ancestral strain.

As anticipated, all SARS-CoV-2 targeting gRNAs tested showed no significant viral suppression against the ancestral nor the D614G strains 1-hour post-infection. The absence of viral suppression with targeting gRNAs at this early timepoint of infection was anticipated because pspCas13b-mediated viral suppression requires intracellular viral replication that had not yet occurred.

In the positive control groups, NCP-1 gRNA is used to target a fully matching sequence within the Nucleocapsid RNA that is conserved in both ancestral and D614G strains. Consistent with the data in **Figure 2** and **Figure 4**, NCP-1 gRNA again showed a very high silencing efficiency, and suppressed 84-88% and 73-74% of replication-competent ancestral and D614G viruses, respectively, at both 24 and 48h timepoints (**Figure 5F & 5H**). The moderate loss of viral suppression in cells infected with the D614G mutant is likely due to the 7-fold higher initial viral load (**Figure 5C, 5E & 5G**). Among the 4 tiled gRNAs targeting the ancestral D614 position in Spike’s RNA, 3 of them (15, 20, and 30) showed efficient viral suppression reaching 78-84% and 44-60% in cells infected with the ancestral and D614G strains respectively, at both 24 and 48h timepoints (**Figure 5F & 5H**). Again, the moderate reduction in D614G suppression with these 3 gRNAs is likely attributable to the initial higher viral load of this D614G strain. Last, the fourth gRNA we tested (harbouring a mismatch with the D614G mutation at spacer position 10) showed the lowest affinity and viral suppression potential with 62-71% suppression of the ancestral strain, and only 3-7% suppression of the D614G, which was not statistically significant at 24 nor at 48h (**Figure 5F & 5H**). This result is congruent with the data obtained from virus-free model (**Figure 5A**), where the gRNA harbouring a single-nucleotide mismatch with D614G at spacer position 10 exhibited the lowest silencing efficiency among the 4 gRNAs tested. This correlation highlights the predictive power of the virus-free model we developed, and emphasises its utility to tractably screen and select the best performing gRNAs. Overall, the data in **Figure 5** provide strong evidence that optimized single gRNAs are likely to retain efficacy against spontaneous point-mutations that arise during viral replication and can efficiently silence ancestral and emerging SARS-CoV-2 mutants, including the D614G.

## DISCUSSION

The remarkable capability of RNA viruses to adapt to selective host and environmental pressure is highly dependent on their ability to generate genomic diversity through the occurrence of *de novo* mutations^34^. Mutation-driven viral evolution can generate drug resistance, immune escape, and potentially increased efficiency of transmission and pathogenicity, all of which are detrimental to the host. Although our understanding of SARS-CoV-2 mutation-driven escape mechanisms remains limited, several reports have demonstrated the emergence of new strains, which possess increased infective potential^8^ or are resistant to monoclonal antibody therapy^7^. In this study, we leveraged a novel technology and employed two key strategies to silence SARS-CoV-2 RNA and counteract its intrinsic ability to escape standard therapies through the generation of *de novo* mutations.

Firstly, the gRNA multiplexing approach to target various regions of viral RNA simultaneously mitigates the risk of target inaccessibility^35^ and potential viral escape through genomic rearrangement or polymorphisms^7,8,14,36^ (**Figure 4, Suppl. Figure 4 & Figure 5**). Given the moderate mutation rate of SARS-CoV-2, it is extremely unlikely that the virus would accumulate simultaneous mutations in various regions to escape a cocktail of 4 gRNAs without compromising its fitness. This multiplexed targeting is comparable to other combination therapeutic strategies that have proven to be effective against other viruses including HIV^37^. Compared to inhibitors of protein function that typically require years of modelling, design, and screening, the advantage of viral RNA targeting with pspCas13b lies in its design-flexibility, predictive efficacy and specificity, and the short time needed to generate the relevant gRNAs. The specificity, high silencing efficiency, and rapid deployment properties of pspCas13b that we have demonstrated are indicative of the translational potential of this technology to reshape the battle against SARS-CoV-2 and provide a means of evolving therapy in concert with viral genomic changes. Likewise, other Cas13 orthologs may also offer similar properties^3,16,19,28,38,39^.

Secondly, our comprehensive mutagenesis analysis revealed that the positively-charged central channel of pspCas13b can tolerate single-nucleotide mismatches within the RNA-RNA hybrid created by spacer-target base-pairing^26,27^ (**Figure 1, 2 & 3**). We demonstrated that a single gRNA can simultaneously silence parental and emerging SARS-CoV-2 strains (**Figure 5**). The mismatch tolerance mechanism shown here revealed that a single gRNA will likely remain effective against future viral mutants that acquire *de novo* single-nucleotide indels as a result of genome replication by the error-prone RdRP of SARS-CoV-2.

Given the high degree of tolerance to single-nucleotide mismatch and the use of gRNA multiplexing, we postulate that a reprogrammed pspCas13b can act as a universal tool to silence various SARS-CoV-2 strains that would escape conventional antiviral therapeutics, including single monoclonal antibodies^7^ or small inhibitor molecules. A key step in enabling clinical translation of this approach will be to develop a safe and effective delivery strategy such as lipid nanoparticle formulations for systemic, and possibly, mucosal delivery^40^. Importantly, a CRISPR-Cas13 based approach is also readily adaptable and expandable to other pathogenic viruses beyond SARS-CoV-2, and may therefore represent a new and powerful approach to antiviral therapeutics.

## Acknowledgements

The authors thanks all lab members from the Trapani, Lewin, Voskoboinik, Subbarao, and Chirlmin Joo labs and VIDRL for facilitating experiments and discussions. This work was supported by the National Health and Medical Research Council (NHMRC) of Australia through a program grant to J.A.T, a program grant and practitioner fellowship to S.R.L. MF is supported by a PeterMac strategic plan funding in partnership with Childhood Cancer Institute Australia (CCIA).

## Data availability

All data are available in the main text and suppl materials. All key plasmids generated in this study will be deposited in Addgene upon publication. The bioinformatic code for the design of single-nucleotide tiled gRNAs and filtering strategies will be made available upon request and will be published in a separate manuscript.

## Authors contribution

M.F conceived the study. M.F, J.A.T and S.R.L supervised the study. M.F, J.A.T, S.R.L, W.Z, W.H, and J.M.L.C designed the experiments and discussed all data. M.F, W.H, and J.M.L.C cloned all constructs, optimized and performed virus-free silencing assays and analysed the data. W.Z performed all SARS-CoV-2 infection assays with help from R.R. W.Z analysed data from infectivity assays. M.F and W.Z extracted viral RNAs, performed RT-PCR, and analysed the data. A.K performed the computational analysis of SARS-CoV-2 guide and target RNAs with input from M.F. I.V, P.G.E, and J.S discussed the project and the data. M.F generated figures and wrote the manuscript. J.A.T and S.R.L revised and edited the manuscript. All authors read, commented, edited, and approved the manuscript.

## METHODS

### Design and cloning of pspCas13b guide RNAs

Individual guides were cloned into the pC0043-PspCas13b^22^ crRNA backbone (addgene#103854, a kind gift from Feng Zhang lab), which we refer to as gRNA backbone. This vector contains pspCas13b gRNA direct repeat (DR) sequence and two BbsI restriction sites for spacer cloning. 20μg of DNA backbone was digested by BbsI (NEB#) following the manufacturer’s instructions (2h at 37C). The digested backbone was gel purified using NucleoSpin™ Gel and PCR Clean-up Kit (Thermo Fisher 12), aliquoted, and stored in −20C°.

For gRNA cloning, a forward and reverse single-stranded DNA oligos containing CACC and CAAC overhangs respectively, were ordered from Sigma (100μM). 1.5μL of 100uM the forward and reverse DNA oligos were annealed in 47μL annealing buffer (5ul NEB buffer 3.1 and 42μL H2O) by 5min incubation at 95C° and slow cooldown in the heating block overnight. 1μL of the annealed oligos were ligated with 0.04ng digested pspCAs13b gRNA backbone in 10μL of T4 ligation buffer (3 hours, RT). All pspCas13b gRNA spacer sequences used in this study are listed in the Table below.

### Cloning of pspCas13b-NES-HIV-T2A-BFP

The original pspCas13b (addgene#103862) is a gift from Feng Zhang lab^22^. A new pspCas13 plasmid was designed by fusing a 3xFlag-T2A-BFP tag to the Cas13 C-terminus using EcoRI and NheI enzymatic restriction. 5μg of pUC19 vector encoding a Psp-Cas13b-3’end-3xFlag-T2A-BFP sequence was generated by DNA synthesis (IDT). Psp-Cas13b-3’end-3xFlag-T2A-BFP and the original pC0046-EF1a-PspCas13b-NES-HIV plasmids were digested by EcoRI/NheI restriction enzymes (1 hour, 37C°), followed by column clean up with NucleoSpin™ Gel and PCR Clean-up Kit (Macherey-Nagel). The digestion efficiency was verified by 1% agarose gel. The Psp-Cas13b-3’end-3xFlag-T2A-BFP fragment with the correct size was gel purified using NucleoSpin™ Gel and PCR Clean-up Kit. The two fragments were ligated using T4 ligase (3 hours, RT), and transformed into Stbl3 chemically competent bacteria. Positive clones were screened by PCR and Singer sequenced (AGRF, AUSTRALIA).

### Cloning of SARS-CoV-2 Spike, Spike D614G, and Nucleocapsid cDNA

A mammalian expression plasmid containing a codon-optimized sequence of the Spike protein was obtained through Genscript plasmid sharing platform (Ref# MC_0101087), and was a kind gift from Dr. Haisheng Yu lab.

The coding sequence of NCP and part of Spike D614G sequence was designed according to the first SARS-CoV-2 genome^23^. The Genscript DNA synthesis platform provided the two sequences that were subsequently cloned into MSCV-IRES-mcherry vector in frame with 3xHA tag using BamHI digestion, gel purification, and ligation with T4 DNA ligase. The ligated product was transformed into chemically competent bacteria (Top10) and positive clones were screened by PCR and Singer sequencing (AGRF, AUSTRALIA).

### Plasmid amplification and purification

TOP10 (for all gRNA, Spike, D614G Spike, and NCP cloning) and Stbl3 (for Cas13 cloning) bacteria were used for transformation. 5-10μL ligated plasmids were transformed into 30 μL of chemically competent bacteria by heat shock at 42C° for 45 seconds, followed by 2min on ice. The transformed bacteria were incubated in 500μL LB broth media containing 75μg/mL ampicillin for 1 hour at 37C° in a shaking incubator (200 rpm). The bacteria were pelleted by centrifugation at 8,000 rpm for 1minute at room temperature (RT), re-suspended in 100 μL of LB broth, and plated onto a prewarmed 10 cm LB agar plate containing 75μg/mL ampicillin, and incubated at 37C° overnight. Next day, single colonies were picked and transferred into bacterial starter cultures and incubated for ~6 hours for mini-prep or maxi-prep DNA purification according to the standard manufacturer’s protocol. All gRNAs and pspCas13b clones that are generated in this study were verified by Sanger sequencing (AGRF, AUSTRALIA).

### Cell culture

HEK293 FT (ATCC CRL-3216) and VERO (ATCC CCL-81) cell lines were cultured in DMEM high glucose media (Life Technologies) containing 10% heat-inactivated fetal bovine serum (Life Technologies), Penicillin/-Streptomycin, and -L-Glutamine (Life Technologies). HEK293 FT and VERO cells were maintained at confluency between 20%-80% in 37C° incubators with either 10% (HEK293 FT) or 5% (VERO). Cells were routinely tested and were mycoplasma negative.

### RNA silencing assays by transient transfection

All transfection experiments were performed in HEK293 FT and cell lines using an optimized Lipofectamine 3000 transfection protocol (Life Technologies, L3000015). For RNA silencing in 293 HEK, cells were plated at approximately 30,000 cells/100μL/96-well in tissue culture treated flat-bottom 96-well plates (Corning) 18 hours prior to transfection. For each well, a total of 100ng DNA plasmids (22ng of pspCas13b-BFP construct, 22ng gRNA plasmid and 56ng of the target gene) were mixed with 0.2μL P3000 reagent in Opti-MEM Serum-free Medium (Life Technologies) to a total of 5μL (mix1). Separately, 4.7μL of Opti-MEM was mixed with 0.3μL Lipofectamine 3000 (mix2). Mix1 and mix2 are added together and incubated for 20 minutes at room temperature, then 10μL of transfection mixture was add to each well. Similar protocol was used for VERO cells transfection except 20,000 cells/well were seeded in a 96-well plate and transfected with a double volume of transfection mixture (20μL). The tables below summarize the transfection protocol used in 96, 24, and 6-well plates for both 293 HEK FT and VERO cells.

| Component (per well) | 96-well | 24-well | 6-well |
| --- | --- | --- | --- |
| 293 HEK FT cells | 30,000 | 150,000 | 750,000 |
| DNA amount | 100ng | 500ng | 2500ng |
| P3000 reagent | 0.2µL | 1µL | 5µL |
| Lipofectamine 3000 reagent | 0.3µL | 1.5µL | 7.5µL |
| Opti-MEM | Up to 10 µL | Up to 50 µL | Up to 250 µL |
| VERO cells | 20,000 | 100,000 | 500,000 |
| DNA amount | 200ng | 1000ng | 5000ng |
| P3000 reagent | 0.4µL | 2µL | 10µL |
| Lipofectamine 3000 reagent | 0.6µL | 3µL | 15µL |
| Opti-MEM | Up to 20 µL | Up to 100 µL | Up to 500 µL |

After transfection, cells were incubated at 37C° and 5% CO2, and the transfection efficiency was monitored by either fluorescence microscopy or FACS analysis.

### Fluorescence microscopy analysis

For RNA silencing experiments, the fluorescence intensity was monitored using EVOS M5000 FL Cell Imaging System (Thermo Fisher). Pictures were taken 48 hours (293 HEK) and 72 hours (VERO) post-transfection, and the fluorescence intensity of each image was quantified using a lab-written macro in ImageJ software. Briefly, all images obtained from a single experiment are simultaneously processed using a batch mode macro. First, images are converted to 8-bit, threshold adjusted, converted to black and white using Convert to Mask function, and fluorescence intensity per pixel measured using Analyze Particles function. Each single mean fluorescence intensity was obtained from four different field of views for each gRNA, and subsequently normalized to the non-targeting (NT) control gRNA. Two-fold or higher reduction in fluorescence intensity is considered as biologically relevant.

### Cell flow cytometry

For monitoring cell transfection/transduction efficacy, cells were re-suspended in 200μL 1x PBS containing 2% FBS for flow cytometry analysis. All samples were analyzed by an LSR II (BD Biosciences), FORTESSA X20 (BD Biosciences) or FACSymphony (BD Biosciences). All flow cytometry profiles were analyzed using FlowJo V10 software (Tree Star Inc).

### Western Blot

Cells were washed three times with ice-cold PBS+/+ and lysed on ice in lysis buffer (50Mm Tris, 150mM NaCl, 1% NP-40, 2% SDS, pH 7.5). Samples were sonicated on low power, two-second pulses, and centrifuged at 16,000g for 10 minutes, 4 °C. Supernatant was transferred to a new tube. Protein concentrations were quantified using the BCA assay (ThermoFisher Scientific) according to the manufacturer’s instructions. 10μg proteins diluted in 1x Bolt LDS sample buffer and 1x Bolt sample reducing agent were denaturated at 95 *°*C for 5 minutes. Samples were resolved by Bolt Bis-Tris Plus 4-12% gels in 1x MES SAS and transferred to PVDF membranes by a Trans-Blot Semi-Dry electrophoretic transfer cell (Bio-Rad) at 20 Volt for 30 minutes. Membranes were incubated in blocking buffer (5% (w/v) skin milk powder in PBST with 0.1% Tween 20) for 1 hour at RT and probed overnight with primary antibody (anti-HA, mAb #2367, Cell Signaling Technology) at 4 °C. Blots were washed three times in PBST with 0.1% Tween20, followed by incubation with HRP-conjugated secondary antibody (Rabbit Anti-Mouse Immunoglobulins/HRP #p0260, Dako) for 1 hour at RT. Membranes were washed in PBST (0.1% Tween20) three times and incubated in appropriate strength ECL detection reagent. Chemiluminescence was detected using Invitrogen iBright Imaging Systems (Thermo Fisher Scientific).

### VERO cells infection assays

VERO cells were transfected in 24-well plates and incubated for 48 h as described above. VERO cells expressing pspCas14b-BFP and various gRNAs were then infected in a level 3 containment laboratory with 200 μL of the ancestral or D614G SARS-CoV-2 virus isolate, a kind gift from Dr. Julian Druce^25^ (Victoria Infectious Disease Reference Lab, VIDRL) at MOI of 0.1 or 0.01 in DMEM containing Penicillin/Streptomycin, L-Glutamine and 1 μg / mL TPCK-treated trypsin (LS003740, Worthington) to facilitate Spike cleavage and cell entry of the virus. After 1-hour incubation at room temperature, Virus production in supernatant was assessed by real-time PCR (RT-PCR) and infectivity assays at various time points.

### RNA extraction, cDNA synthesis and RT-PCR

Viral RNA from 140 μL cell culture supernatant was extracted using the QIAamp Viral RNA Mini kit (#52906, Qiagen) following the manufacturer’s instructions. SARS-CoV-2 RNA was converted to cDNA using the SensiFAST cDNA kit (#BIO-65053, BioLine) with 10 μL of RNA extract per reaction following the manufacturer’s instructions. Quantitative RT-PCR reaction targeting RdRP gene was performed in triplicate in a Mx3005P QPCR System (Agilent) using PrecisionFAST qPCR Master Mix (PFAST-LR-1, Integrated Science). Total reaction mixture contains 2.5 μL cDNA, 0.75 μM forward primer (5’-AAA TTC TAT GGT GGT TGG CAC AAC ATG TT-3’, 0.75 μM reverse primer (5’-TAG GCA TAG CTC TRT CAC AYT T-3’) and 0.15 μM TaqMan Probe (5’-FAM-TGG GTT GGG ATT ATC-MGBNFQ-3’).

### Infectivity assays

For titration of the 50% tissue culture infectious dose (TCID50) of SARS-CoV-2, VERO cells were plated in 96-well plates at 20,000 cells per well in MEM containing 5% heat-inactivated fetal bovine serum, Penicillin/Streptomycin, L-Glutamine and 15 mM HEPES (Life Technologies). The cells were incubated overnight in a 5% CO2 environment at 37°C, washed once with PBS and then cultured in serum-free MEM containing Penicillin/Streptomycin, L-Glutamine, 15 mM HEPES and 1 μg / mL TPCK-treated trypsin. A 10-fold initial dilution of samples with one freeze-thaw cycle was made in quadruplicate wells of the 96-well plates followed by 6 serial 10-fold dilutions. The last row served as negative control without addition of any sample. After a 4-day incubation, the plates were observed for the presence of cytopathogenic effect (CPE) using an inverted optical microscope. Any sign of CPE was categorized as positive results. The endpoint titres were calculated by means of a simplified Reed & Muench method^41^.

### Data analysis

Data analyses and visualization (graphs) were performed in GraphPad Prism software version 7. Specific statistical tests, numbers of biological replicates are mentioned in respective figure legends. The silencing efficiency of various gRNAs was analyzed using one-way Anova followed by Dunnett’s multiple comparison test where we compare every mean to a control mean as indicated in the figures (95% confidence interval). Unpaired Student’s t-test (95% confidence interval) was used to compare the viral titre between ancestral and D614G. The P values (P) are indicated in the figures. P<0.05 is considered as statistically significant.

### Spacer sequence of gRNAs used in this study

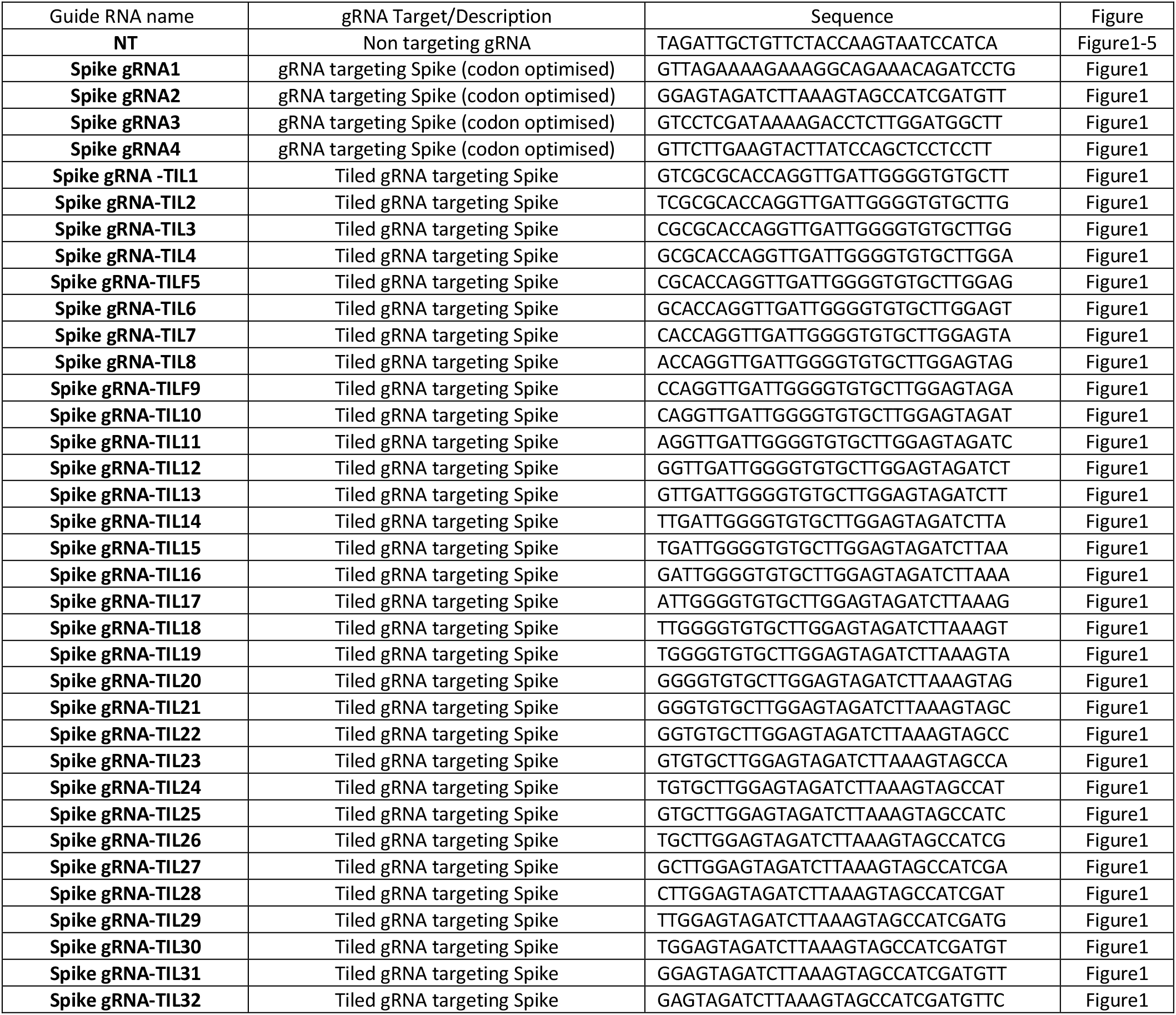
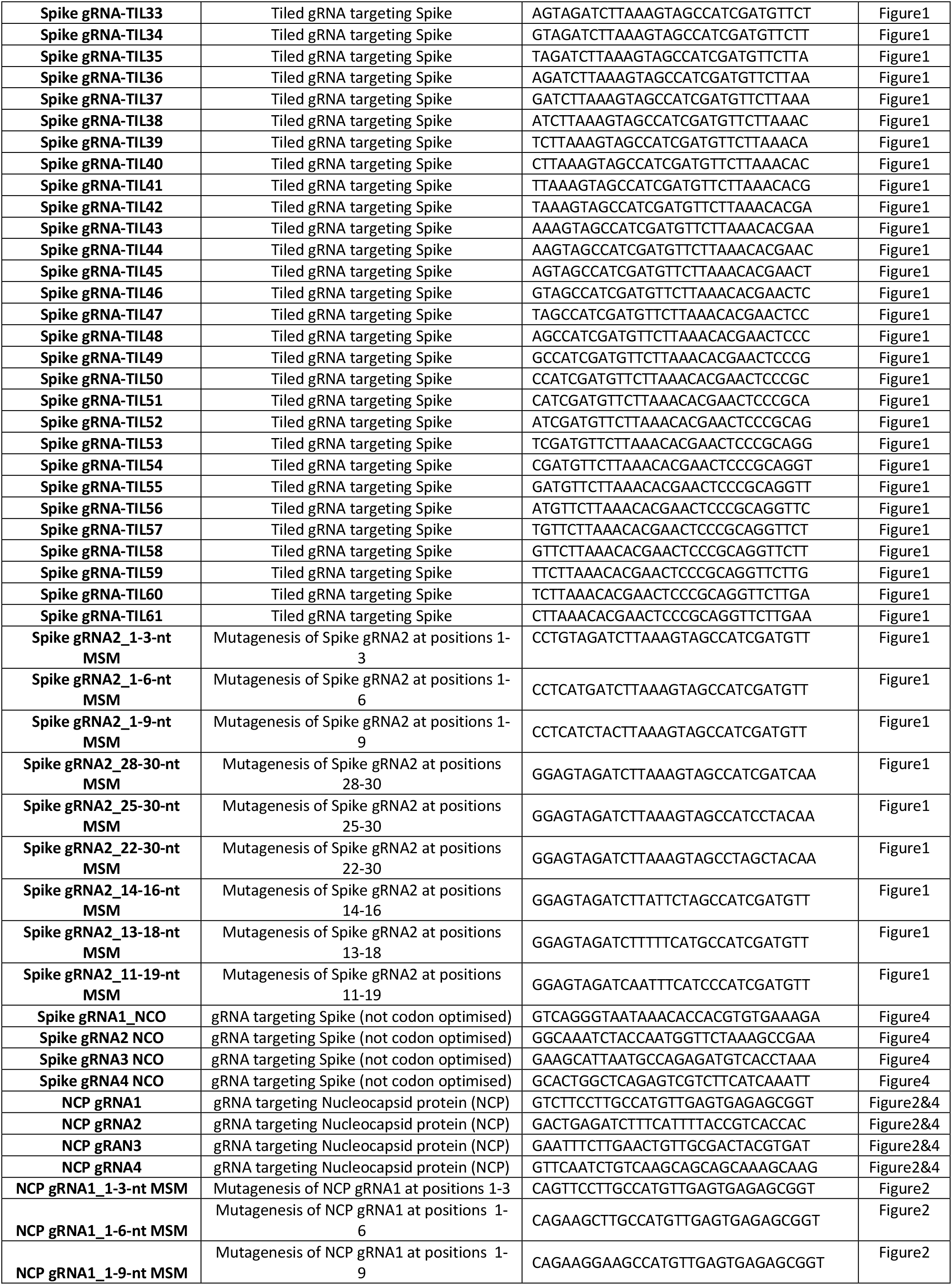
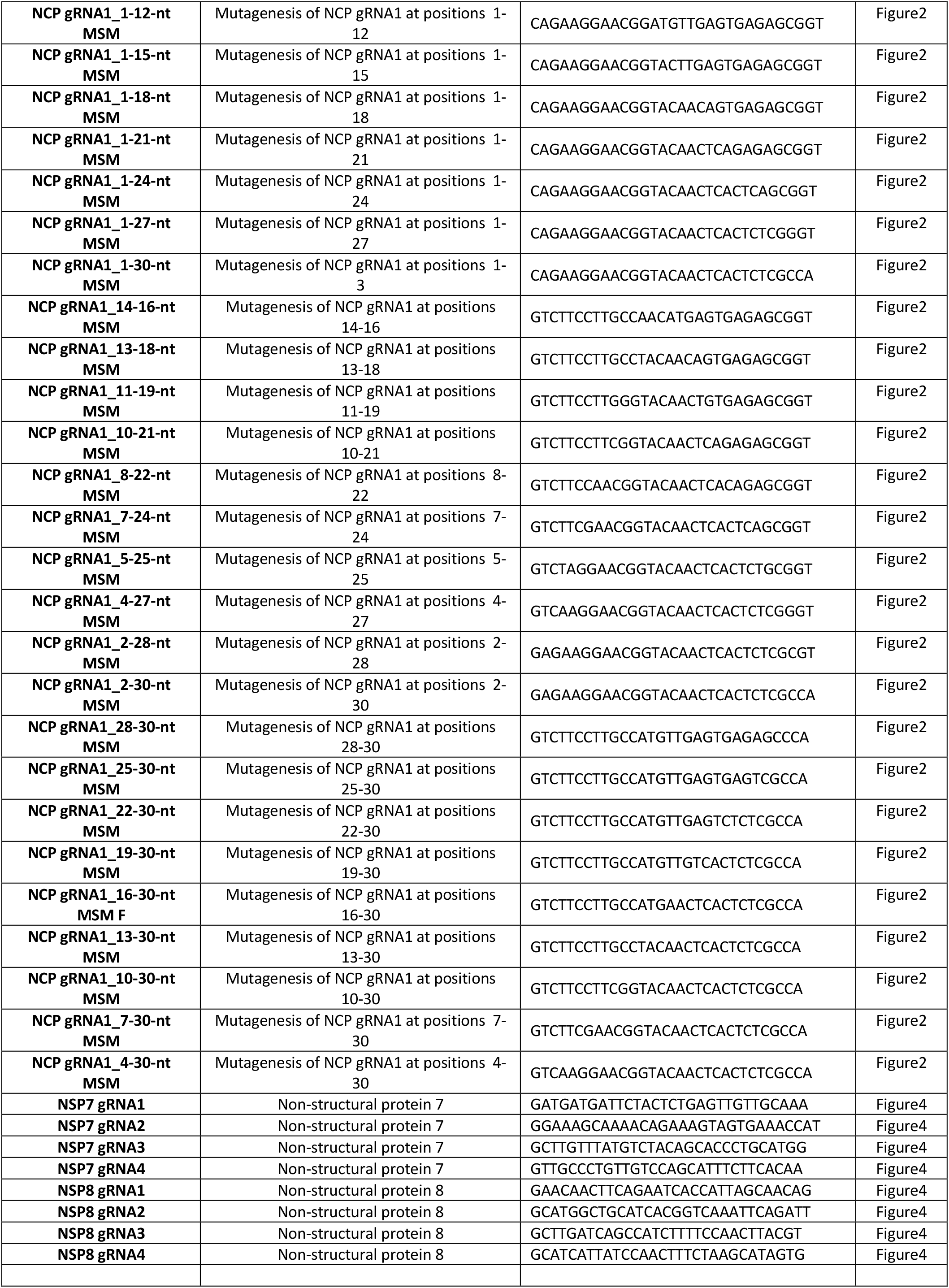
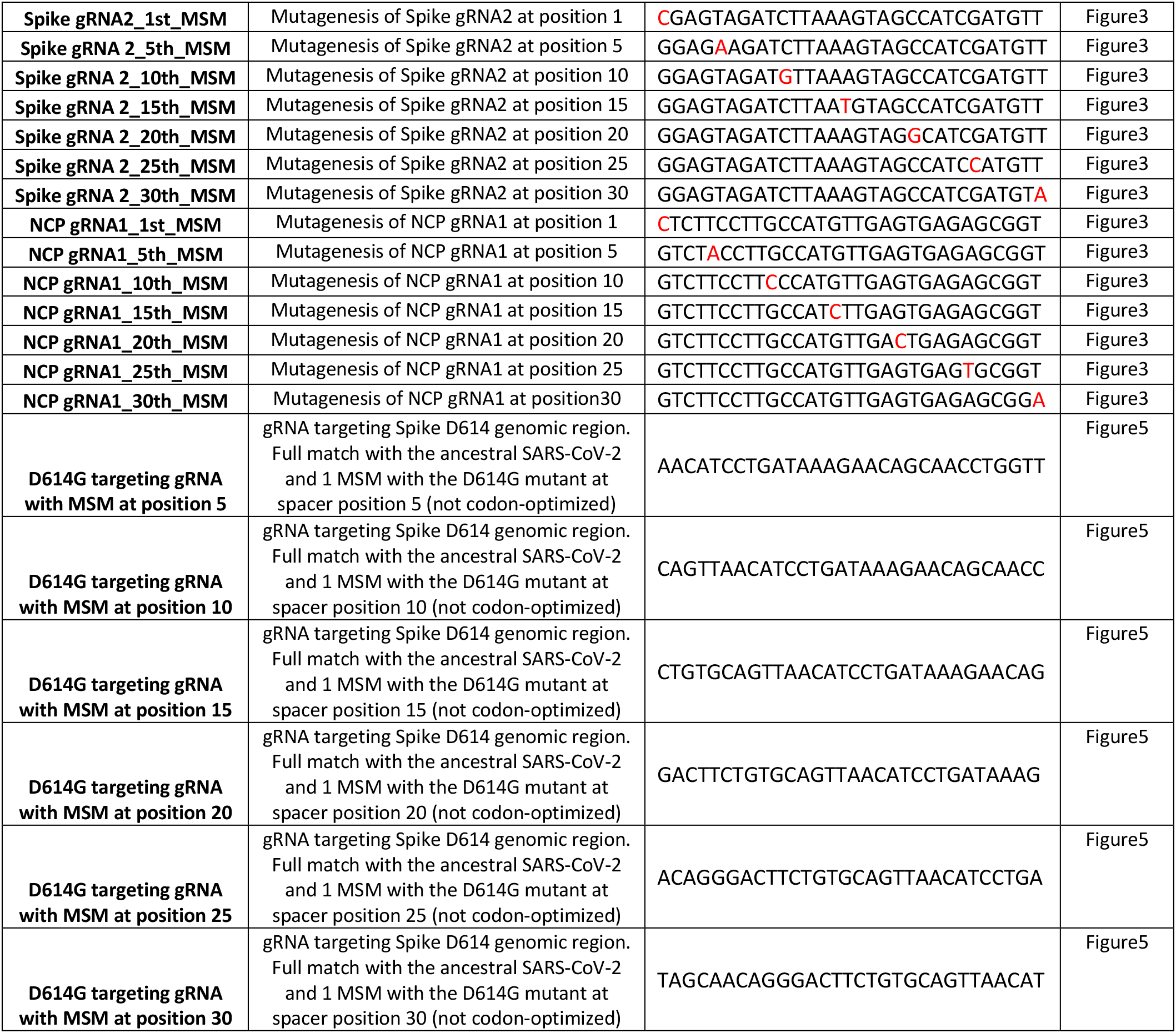

### cDNA sequences of constructs used in this study

#### PspCasl3b-3xFLAG-T2A-BFP

atgaacatccccgctctggtggaaaaccagaagaagtactttggcacctacagcgtgatggccatgctgaacgctcagaccgtgctggaccacatccagaaggtggccgatattgagggcgagcagaacgagaacaacgagaatctgtggtttcaccccgtgatgagccacctgtacaacgccaagaacggctacgacaagcagcccgagaaaaccatgttcatcatcgagcggctgcagagctacttcccattcctgaagatcatggccgagaaccagagagagtacagcaacggcaagtacaagcagaaccgcgtggaagtgaacagcaacgacatcttcgaggtgctgaagcgcgccttcggcgtgctgaagatgtacagggacctgaccaaccactacaagacctacgaggaaaagctgaacgacggctgcgagttcctgaccagcacagagcaacctctgagcggcatgatcaacaactactacacagtggccctgcggaacatgaacgagagatacggctacaagacagaggacctggccttcatccaggacaagcggttcaagttcgtgaaggacgcctacggcaagaaaaagtcccaagtgaataccggattcttcctgagcctgcaggactacaacggcgacacacagaagaagctgcacctgagcggagtgggaatcgccctgctgatctgcctgttcctggacaagcagtacatcaacatctttctgagcaggctgcccatcttctccagctacaatgcccagagcgaggaacggcggatcatcatcagatccttcggcatcaacagcatcaagctgcccaaggaccggatccacagcgagaagtccaacaagagcgtggccatggatatgctcaacgaagtgaagcggtgccccgacgagctgttcacaacactgtctgccgagaagcagtcccggttcagaatcatcagcgacgaccacaatgaagtgctgatgaagcggagcagcgacagattcgtgcctctgctgctgc agtatatcgattacggcaagctgttcgaccacatcaggttccacgtgaacatgggcaagctgagatacctgctgaaggccgacaagacctgcatcgacggccagaccagagtcagagtgatcgagcagcccctgaacggcttcggcagactggaagaggccgagacaatgcggaagcaagagaacggcaccttcggcaacagcggcatccggatcagagacttcgagaacatgaagcgggacgacgccaatcctgccaactatccctacatcgtggacacctacacacactacatcctggaaaacaacaaggtcgagatgtttatcaacgacaaagaggacagcgccccactgctgcccgtgatcgaggatgatagatacgtggtcaagacaatccccagctgccggatgagcaccctggaaattccagccatggccttccacatgtttctgttcggcagcaagaaaaccgagaagctgatcgtggacgtgcacaaccggtacaagagactgttccaggccatgcagaaagaagaagtgaccgccgagaatatcgccagcttcggaatcgccgagagcgacctgcctcagaagatcctggatctgatcagcggcaatgcccacggcaaggatgtggacgccttcatcagactgaccgtggacgacatgctgaccgacaccgagcggagaatcaagagattcaaggacgaccggaagtccattcggagcgccgacaacaagatgggaaagagaggcttcaagcagatctccacaggcaagctggccgacttcctggccaaggacatcgtgctgtttcagcccagcgtgaacgatggcgagaacaagatcaccggcctgaactaccggatcatgcagagcgccattgccgtgtacgatagcggcgacgattacgaggccaagcagcagttcaagctgatgttcgagaaggcccggctgatcggcaagggcacaacagagcctcatccatttctgtacaaggtgttcgcccgcagcatccccgccaatgccgtcgagttctacgagcgctacctgatcgagcggaagttctacctgaccggcctgtccaacgagatcaagaaaggcaacagagtggatgtgcccttcatccggcgggaccagaacaagtggaaaacacccgccatgaagaccctgggcagaatctacagcgaggatctgcccgtggaactgcccagacagatgttcgacaatgagatcaagtcccacctgaagtccctgccacagatggaaggcatcgacttcaacaatgccaacgtgacctatctgatcgccgagtacatgaagagagtgctggacgacgacttccagaccttctaccagtggaaccgcaactaccggtacatggacatgcttaagggcgagtacgacagaaagggctccctgcagcactgcttcaccagcgtggaagagagagaaggcctctggaaagagcgggcctccagaacagagcggtacagaaagcaggccagcaacaagatccgcagcaaccggcagatgagaaacgccagcagcgaagagatcgagacaatcctggataagcggctgagcaacagccggaacgagtaccagaaaagcgagaaagtgatccggcgctacagagtgcaggatgccctgctgtttctgctggccaaaaagaccctgaccgaactggccgatttcgacggcgagaggttcaaactgaaagaaatcatgcccgacgccgagaagggaatcctgagcgagatcatgcccatgagcttcaccttcgagaaaggcggcaagaagtacaccatcaccagcgagggcatgaagctgaagaactacggcgacttctttgtgctggctagcgacaagaggatcggcaacctgctggaactcgtgggcagcgacatcgtgtccaaagaggatatcatggaagagttcaacaaatacgaccagtgcaggcccgagatcagctccatcgtgttcaacctggaaaagtgggccttcgacacataccccgagctgtctgccagagtggaccgggaagagaaggtggacttcaagagcatcctgaaaatcctgctgaacaacaagaacatcaacaaagagcagagcgacatcctgcggaagatccggaacgccttcgatcacaacaattaccccgacaaaggcgtggtggaaatcaaggccctgcctgagatcgccatgagcatcaagaaggcctttggggagtacgccatcatgaagggatcccttcaactgcctccacttgaaagactgacactggactataaggaccacgacggagactacaaggatcatgatattgattacaaagacgatgacgataagggcggcgggtccggaggagagggcagaggaagtctcctaacatgcggtgacgtggaggagaatcctggcccaatgagcgagctgattaaggagaacatgcacatgaagctgtacatggagggcaccgtggacaaccatcacttcaagtgcacatccgagggcgaaggcaagccctacgagggcacccagaccatgagaatcaaggtggtcgagggcggccctctccccttcgccttcgacatcctggctactagcttcctctacggcagcaagaccttcatcaaccacacccagggcatccccgacttcttcaagcagtccttccctgagggcttcacatgggagagagtcaccacatacgaggacgggggcgtgctgaccgctacccaggacaccagcctccaggacggctgcctcatctacaacgtcaagatcagaggggtgaacttcacatccaacggccctgtgatgcagaagaaaacactcggctgggaggccttcaccgagaccctgtaccccgctgacggcggcctggaaggcagaaacgacatggccctgaagctcgtgggcgggagccatctgatcgcaaacatcaagaccacatatagatccaagaaacccgctaagaacctcaagatgcctggcgtctactatgtggactacagactggaaagaatcaaggaggccaacaacgagacctacgtcgagcagcacgaggtggcagtggccagatactgcgacctccctagcaaactggggcacaagcttaattga

Black: PspCas13b

Green:3xFlag

Red: T2A tag

Blue: BFP

#### Coding sequence of codon optimized Spike protein-P2A-EGFP

atgttcgtgtttctggtgctgctgcctctggtgagctcccagtgcgtgaacctgaccacacggacacagctgccccctgcctacaccaacagcttcacaaggggcgtgtactaccccgacaaggtgtttagatctagcgtgctgcactccacacaggatctgtttctgcctttcttttctaacgtgacctggttccacgctatccacgtgtccggcaccaacggaacaaagaggttcgacaacccagtgctgccctttaacgat ggcgtgtacttcgcctccaccgagaagtctaacatcatcagaggctggatctttggaaccacactggacagcaagacacagtccctgctgatcgtgaacaacgccaccaacgtggtcatcaaggtgtgcgagttccagttttgtaacgatccattcctgggcgtgtactaccacaagaacaacaagtcttggatggagagcgagtttcgcgtgtactcctctgccaacaactgtacatttgagtacgtgtcccagcccttcctgatggacctggagggcaagcagggaaacttcaagaacctgcgggagttcgtgtttaagaacatcgatggctactttaagatctactccaagcacaccccaatcaacctggtgcgcgacctgccacagggcttctctgccctggagccactggtggatctgcccatcggaatcaacatcaccaggtttcagacactgctggccctgcacagaagctacctgacaccaggcgacagctcctctggatggaccgctggagctgctgcctactacgtgggctacctgcagccccggaccttcctgctgaagtacaacgagaacggaaccatcacagacgctgtggattgcgccctggaccccctgtctgagaccaagtgtacactgaagagctttaccgtggagaagggcatctaccagacaagcaacttccgggtgcagcctaccgagtccatcgtgcgctttcccaacatcacaaacctgtgcccttttggagaggtgttcaacgctacccgcttcgcctccgtgtacgcttggaaccggaagcgcatctccaactgcgtggccgactactctgtgctgtacaacagcgccagcttcagcaccttcaagtgctacggcgtgagcccaacaaagctgaacgacctgtgctttaccaacgtgtacgctgattccttcgtgatcaggggagacgaggtgcgccagatcgctcccggccagacaggaaagatcgctgactacaactacaagctgcctgacgatttcaccggctgcgtgatcgcctggaactctaacaacctggatagcaaagtgggcggaaactacaactacctgtacaggctgtttagaaagtctaacctgaagccattcgagcgggacatctccacagagatctaccaggctggctctaccccatgcaacggagtggagggcttcaactgttacttccctctgcagagctacggattccagccaacaaacggcgtgggataccagccctaccgcgtggtggtgctgtcttttgagctgctgcacgctcctgctacagtgtgcggaccaaagaagagcaccaacctggtgaagaacaagtgcgtgaacttcaactttaacggactgaccggcacaggagtgctgaccgagtctaacaagaagttcctgccttttcagcagttcggccgggacatcgccgataccacagacgctgtgcgcgaccctcagaccctggagatcctggatatcacaccatgctccttcggcggagtgtctgtgatcacaccaggaaccaacacaagcaaccaggtggccgtgctgtaccaggacgtgaactgtaccgaggtgcccgtggctatccacgccgatcagctgacccctacatggagggtgtactctaccggcagcaacgtgttccagacaagagccggctgtctgatcggagctgagcacgtgaacaacagctacgagtgcgacatccctatcggcgccggaatctgtgcttcctaccagacccagacaaactccccaaggagagccaggtctgtggctagccagtccatcatcgcctacaccatgagcctgggcgccgagaactccgtggcttactccaacaactctatcgctatccctaccaacttcacaatctccgtgaccacagagatcctgccagtgagcatgaccaagacatccgtggactgcacaatgtacatctgtggagattccaccgagtgctctaacctgctgctgcagtacggctctttctgtacccagctgaacagagccctgacaggaatcgctgtggagcaggacaagaacacacaggaggtgttcgcccaggtgaagcagatctacaagaccccacccatcaaggactttggcggattcaactttagccagatcctgcccgatcctagcaagccatccaagaggtcttttatcgaggacctgctgttcaacaaggtgaccctggctgatgccggcttcatcaagcagtacggcgattgcctgggagacatcgctgccagagacctgatctgtgcccagaagtttaacggactgaccgtgctgcctccactgctgacagatgagatgatcgctcagtacacatctgctctgctggccggcaccatcacaagcggatggaccttcggcgctggagctgccctgcagatcccctttgccatgcagatggcttacagattcaacggcatcggagtgacccagaacgtgctgtacgagaaccagaagctgatcgccaaccagtttaactccgctatcggcaagatccaggactctctgagctccacagctagcgccctgggaaagctgcaggatgtggtgaaccagaacgctcaggccctgaacaccctggtgaagcagctgtctagcaacttcggcgccatctcctctgtgctgaacgatatcctgagcaggctggacaaggtggaggctgaggtgcagatcgacaggctgatcacaggaagactgcagtccctgcagacctacgtgacacagcagctgatcagggctgctgagatcagggcttctgccaacctggctgccaccaagatgagcgagtgcgtgctgggccagtccaagagagtggacttttgtggcaagggataccacctgatgagcttcccacagtccgcccctcacggagtggtgtttctgcacgtgacctacgtgccagctcaggagaagaacttcaccacagctcccgccatctgccacgatggcaaggcccactttcctcgggagggcgtgttcgtgagcaacggaacccactggtttgtgacacagcgcaacttctacgagccacagatcatcaccacagacaacacattcgtgtccggcaactgtgacgtggtcatcggaatcgtgaacaacaccgtgtacgatcctctgcagccagagctggactcttttaaggaggagctggataagtacttcaagaaccacaccagccctgacgtggatctgggcgacatctctggaatcaacgccagcgtggtgaacatccagaaggagatcgaccggctgaacgaggtggctaagaacctgaacgagtccctgatcgatctgcaggagctgggcaagtacgagcagtacatcaagtggccctggtacatctggctgggcttcatcgccggactgatcgctatcgtgatggtgaccatcatgctgtgctgtatgacaagctgctgttcctgcctgaagggctgctgttcttgtggaagctgctgtaagtttgacgaggacgatagcgagcctgtgctgaagggcgtgaagctgcactacaccaagcttggatccggaagcggagctactaacttcagcctgctgaagcaggctggagacgtggaggagaaccctggacctatgagcaagggcgaggagctgttcaccggggtggtgcccatcctggtcgagctggacggcgacgtaaacggccacaagttcagcgtgtccggcgagggcgagggcgatgccacctacggcaagctgaccctgaagttcatctgcaccaccggcaagctgcccgtgccctggcccaccctcgtgaccaccctgacctacggcgtgcagtgcttcagccgctaccccgaccacatgaagcagcacgacttcttcaagtccgccatgcccgaaggctacgtccaggagcgcaccatcttcttcaaggacgacggcaactacaagacccgcgccgaggtgaagttcgagggcgacaccctggtgaaccgcatcgagctgaagggcatcgacttcaaggaggacggcaacatcctggggcacaagctggagtacaacta caacagccacaacgtctatatcatggccgacaagcagaagaacggcatcaaggtgaacttcaagatccgccacaacatcgaggacggcagcgtgcagctcgccgaccactaccagcagaacacccccatcggcgacggccccgtgctgctgcccgacaaccactacctgagcacccagtccgccctgagcaaagaccccaacgagaagcgcgatcacatggtcctgctggagttcgtgaccgccgccgggatcactcacggcatggacgagctgtacaagtaa

Blake: Codon-optimized Spike sequence

Blue: P2A tag

Green: EGFP

#### Coding sequence of Nucleocapsid protein-3xHA

atgtctgataatggaccccaaaatcagcgaaatgcaccccgcattacgtttggtggaccctcagattcaactggcagtaaccagaatggagaacgcagtggggcgcgatcaaaacaacgtcggccccaaggtttacccaataatactgcgtcttggttcaccgctctcactcaacatggcaaggaagaccttaaattccctcgaggacaaggcgttccaattaacaccaatagcagtccagatgaccaaattggctactaccgaagagctaccagacgaattcgtggtggtgacggtaaaatgaaagatctcagtccaagatggtatttctactacctaggaactgggccagaagctggacttccctatggtgctaacaaagacggcatcatatgggttgcaactgagggagccttgaatacaccaaaagatcacattggcacccgcaatcctgctaacaatgctgcaatcgtgctacaacttcctcaaggaacaacattgccaaaaggcttctacgcagaagggagcagaggcggcagtcaagcctcttctcgttcctcatcacgtagtcgcaacagttcaagaaattcaactccaggcagcagtaggggaacttctcctgctagaatggctggcaatggcggtgatgctgctcttgctttgctgctgcttgacagattgaaccagcttgagagcaaaatgtctggtaaaggccaacaacaacaaggccaaactgtcactaagaaatctgctgctgaggcttctaagaagcctcggcaaaaacgtactgccactaaagcatacaatgtaacacaagctttcggcagacgtggtccagaacaaacccaaggaaattttggggaccaggaactaatcagacaaggaactgattacaaacattggccgcaaattgcacaatttgcccccagcgcttcagcgttcttcggaatgtcgcgcattggcatggaagtcacaccttcgggaacgtggttgacctacacaggtgccatcaaattggatgacaaagatccaaatttcaaagatcaagtcattttgctgaataagcatattgacgcatacaaaacattcccaccaacagagcctaaaaaggacaaaaagaagaaggctgatgaaactcaagccttaccgcagagacagaagaaacagcaaactgtgactcttcttcctgctgcagatttggatgatttctccaaacaattgcaacaatccatgagcagtgctgactcaactcaggccggtagttcctacccatacgatgttccagattacgcttatccctacgacgtgcctgattatgcatacccatatgatgtccccgactatgcctaa

Black: Nucleocapsid protein

Blue: 3xHA tag

#### Coding sequence of Spike D614G-3XHA

atgtttgtaattagaggtgatgaagtcagacaaatcgctccagggcaaactggaaagattgctgattataattataaattaccagatgattttacaggctgcgttatagcttggaattctaacaatcttgattctaaggttggtggtaattataattacctgtatagattgtttaggaagtctaatctcaaaccttttgagagagatatttcaactgaaatctatcaggccggtagcacaccttgtaatggtgttgaaggttttaattgttactttcctttacaatcatatggtttccaacccactaatggtgttggttaccaaccatacagagtagtagtactttcttttgaacttctacatgcaccagcaactgtttgtggacctaaaaagtctactaatttggttaaaaacaaatgtgtcaatttcaacttcaatggtttaacaggcacaggtgttcttactgagtctaacaaaaagtttctgcctttccaacaatttggcagagacattgctgacactactgatgctgtccgtgatccacagacacttgagattcttgacattacaccatgttcttttggtggtgtcagtgttataacaccaggaacaaatacttctaaccaggttgctgttctttatcagggtgttaactgcacagaagtccctgttgctattcatgcagatcaacttactcctacttggcgtgtttattctacaggttctaatgtttttcaaacacgtgcaggctgtttaataggggctgaacatgtcaacaactcatatgagtgtgacatacccattggtgcaggtatatgcgctagttatcagactcagactaattctcctcggcgggcacgtagtgtagctagtcaatccatcattgcctacactatgtcacttggtgcagaaaattcagttgcttactctaataactctattgccatacccacaaattttactattagtgttaccacagaaattctaccagtgtctatgaccaagacatcagtagattgtacaatgtacatttgtggtgattcaactgaatgcagcaatcttttgttgcaatatggcagtttttgtacacaattaaaccgtgctttaactggaatagctgttgaacaagacaaaaacacccaagaagtttttgcacaagtcaaacaaatttacaaaacaccaccaattaaagattttggtggttttaatttttcacaaatattggtagttcctacccatacgatgttccagattacgcttatccctacgacgtgcctgattatgcatacccatatgatgtccccgactatgcctaa

Black: part of the Spike sequence

Blue: 3xHa tag

## Supplementary Figures legends

**Supplementary Table 1.** Single-nucleotide increment spacer sequence (DNA) of gRNAs covering the entire genome of SARS-CoV-2. The 30-nt spacer sequences were generated using inhouse script written in Python and are complementary to the target RNA. For each gRNA, we provide the matching location in the SARS-CoV-2 genome, the predicted secondary structure of the spacer and target sequences, and the corresponding predicted minimum free energy (kcal/mol). The data in this table are unfiltered and unranked.

**Supplementary Table 2.** Single-nucleotide increment spacer sequence of gRNAs covering the entire genome of SARS-CoV-2 are listed. The 30-nt spacer sequences are complementary to the target, and were generated using inhouse code written in Python. For each gRNA, we provide the matching coordinate in the SARS-CoV-2 genome, the predicted secondary structure of the spacer and target sequences, and the predicted minimum free energy (kcal/mol). The data in this table are filtered to remove spacer sequences that include 4 or more successive T residues that would prematurely terminate Pol III-driven spacer transcription and generate non-functional gRNAs. The gRNAs are ranked based on predicted secondary structures in the target and spacer sequence.

**Supplementary Table 3.** Top scoring 839 gRNAs with a predicted open secondary structure in the spacer and target sequences. These gRNAs are designed to fully base-pair with various regions of SRAS-CoV-2 genome and are predicted to achieve high silencing efficiency, given that Cas13b efficacy is governed by target accessibility.

**Supplementary Table 4.** All gRNAs used in this study are listed.

**Supplementary Figure 1.**
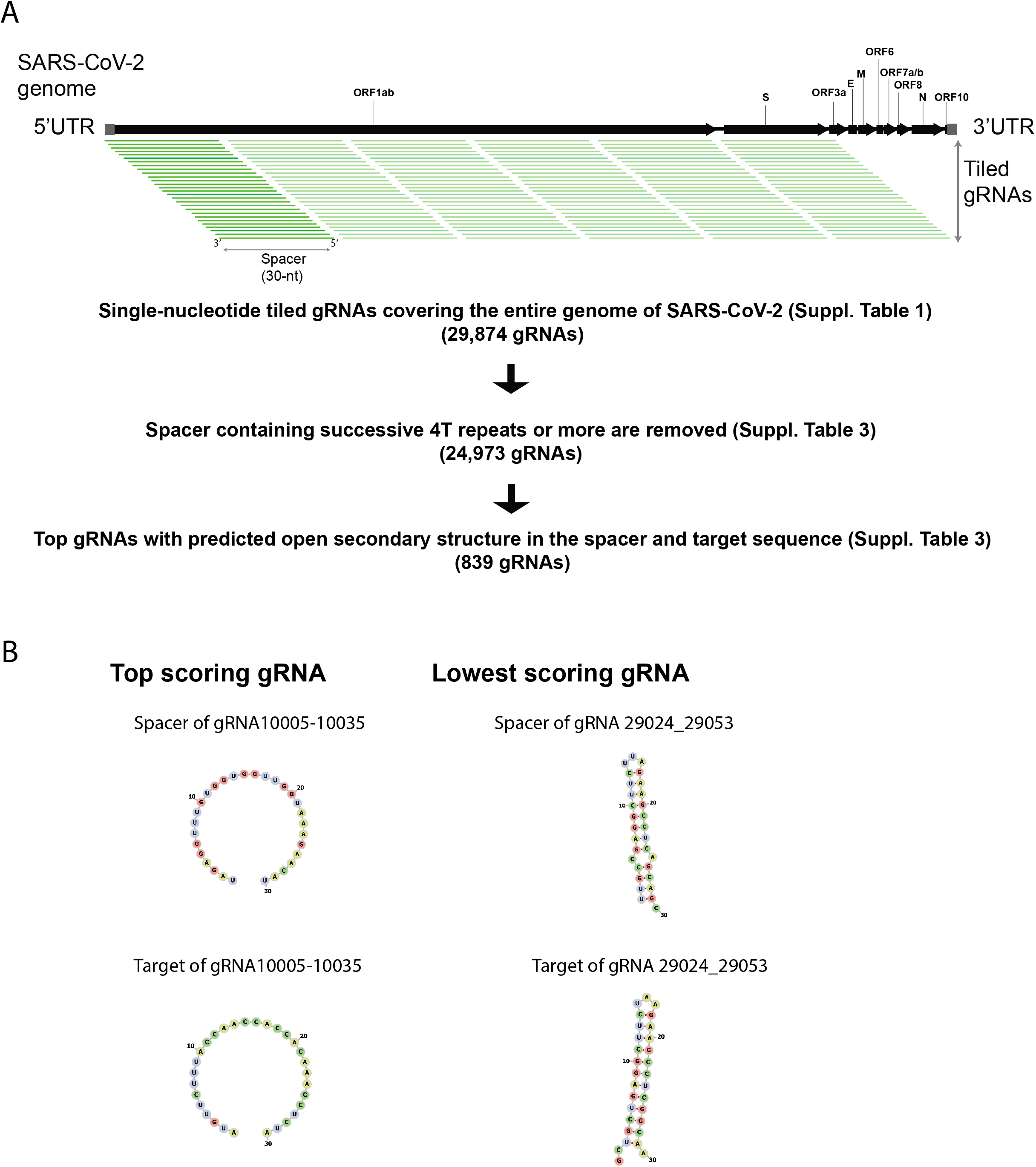
**(A)** Bioinformatics pipeline for the design and selection of pspCas13b gRNAs that are predicted to achieve high silencing efficiency. gRNAs are tiled across the entire RNA genome of SARS-CoV-2 with single-nucleotide increments. gRNAs harbouring 4 or more successive T residues are removed to avoid premature termination of Pol III-driven gRNA transcription, and the top 839 gRNAs with predicted open secondary structure in the spacer and target are selected. **(B)** Predicted secondary structure of the spacer (upper panels) and its target sequence (lower panels) in the top scoring and lowest scoring gRNAs in Suppl. Table 1. The RNA secondary structures were generated using the RNAfold program^1^ (ViennaRNA webservices). The top scoring gRNA shows a predicted open structure for both the spacer and the target, while both structures of the lowest scoring gRNA exhibit high probability of folding into stem-loop that may impair gRNA loading into pspCas13b and target accessibility.

**Supplementary Figure 2.**
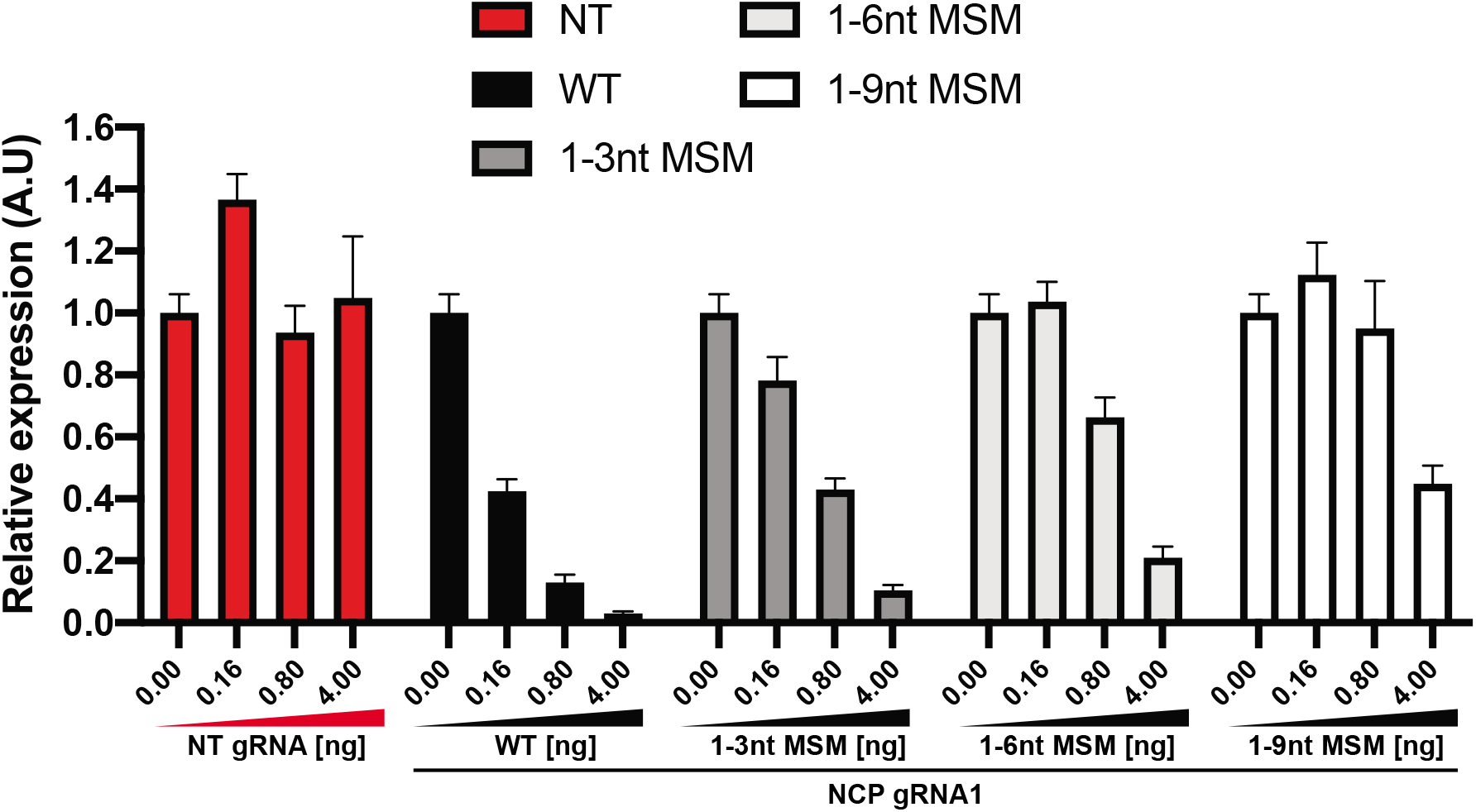
Dose-dependent silencing of NCP transcript with nontargeting gRNA (NT), wildtype gRNA1 (WT), gRNA1 harbouring 1-3, 1-6, or 1-9 nucleotides mismatches at the 5’end of the spacer. Data are normalized mean fluorescence and errors are SD from 4 different field of views. N=1. The dose-dependent silencing shows that the silencing efficiency is dependent on the degree of base-pairing between the spacer and the target. The reduction in the silencing efficiency is proportional to the number of mismatches within the spacer-target RNA-RNA duplex. For instance, with 0.8ng of WT, 1-3, 1-6, and 1-9 nucleotides mismatches gRNAs we achieved 87%, 57%, 34%, and 5% silencing efficiency, respectively. Thus, demonstrating the dependency between the spacer-target basepairing and silencing efficiency.

**Supplementary Figure 3.**
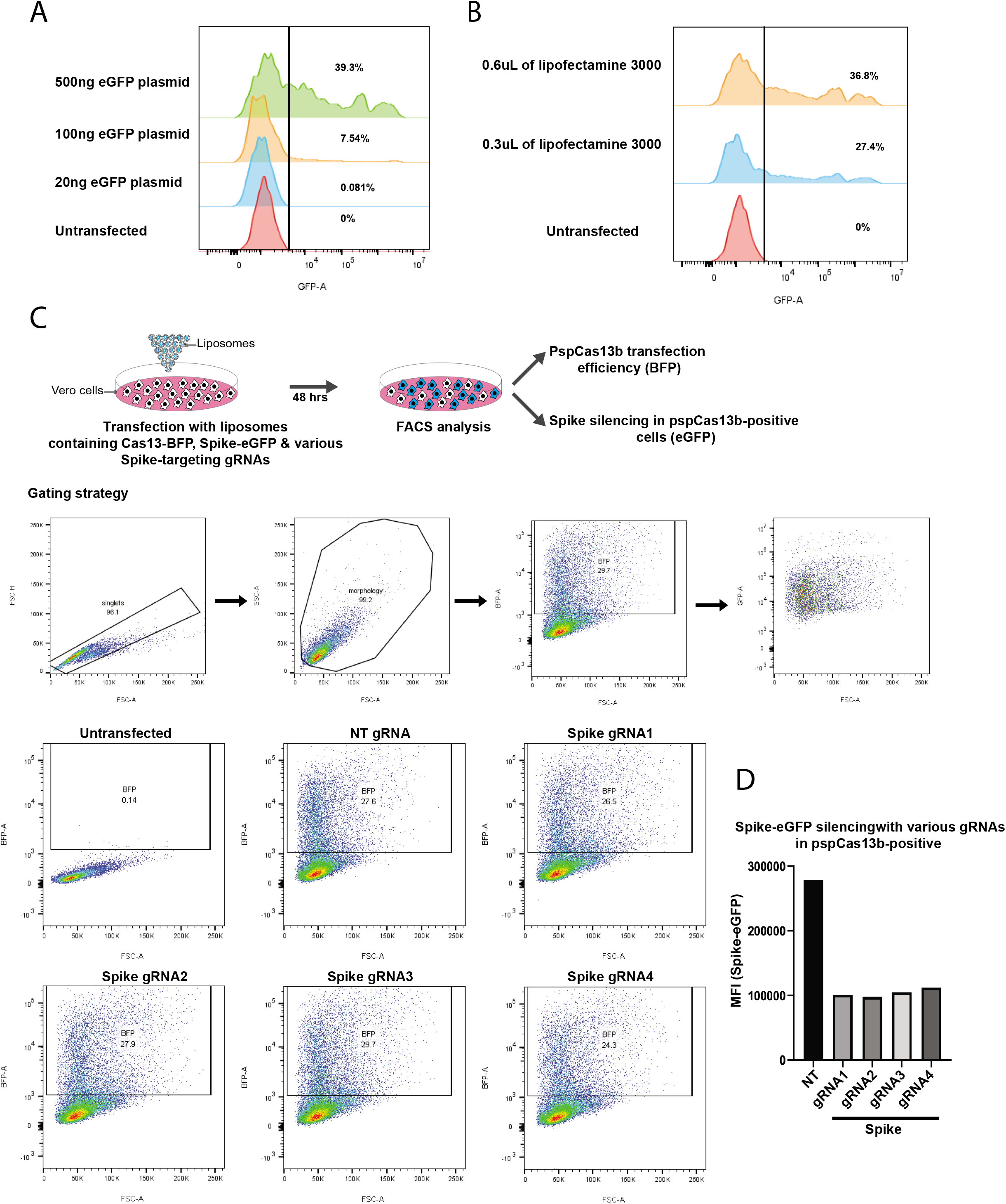
Optimization of VERO cells transfection conditions. (**A**) 20,000 VERO cells were seeded in a 96-well plate and transfected with various amounts of Spike-eGFP reporter plasmid (0-500ng). 48h later, the transfection efficiency was measured by flow cytometry (FACS). (**B**) 20,000 VERO cells were seeded in a 96-well plate and transfected with 100ng of Spike-eGFP reporter plasmid packaged in either 0.3μL or 0.6μL of lipofectamine 3000. Flow cytometry analysis showed that 0.6μL of lipofectamine 3000 gave the highest transfection efficiency (36.8%). N=1. (**C**) FACS analysis of pspCas13b-BFP transfection efficiency in VERO cells. VERO cells were transfected with pspCas13b-BFP, Spike-eGFP, and either NT or four Spike-targeting gRNAs. Both transfection and silencing efficiencies were measured by FACS 48h post-transfection. The gating strategy is shown in the upper panels. In this experiment 25-30% of VERO cells expressed detectable levels of pspCas13b-BFP. (**D**) The histogram quantifies eGFP mean fluorescence intensity (MFI) in each transfection condition that reflects the silencing of Spike-eGFP RNA in pspCas13b-positive VERO cells.

**Supplementary Figure 4.**
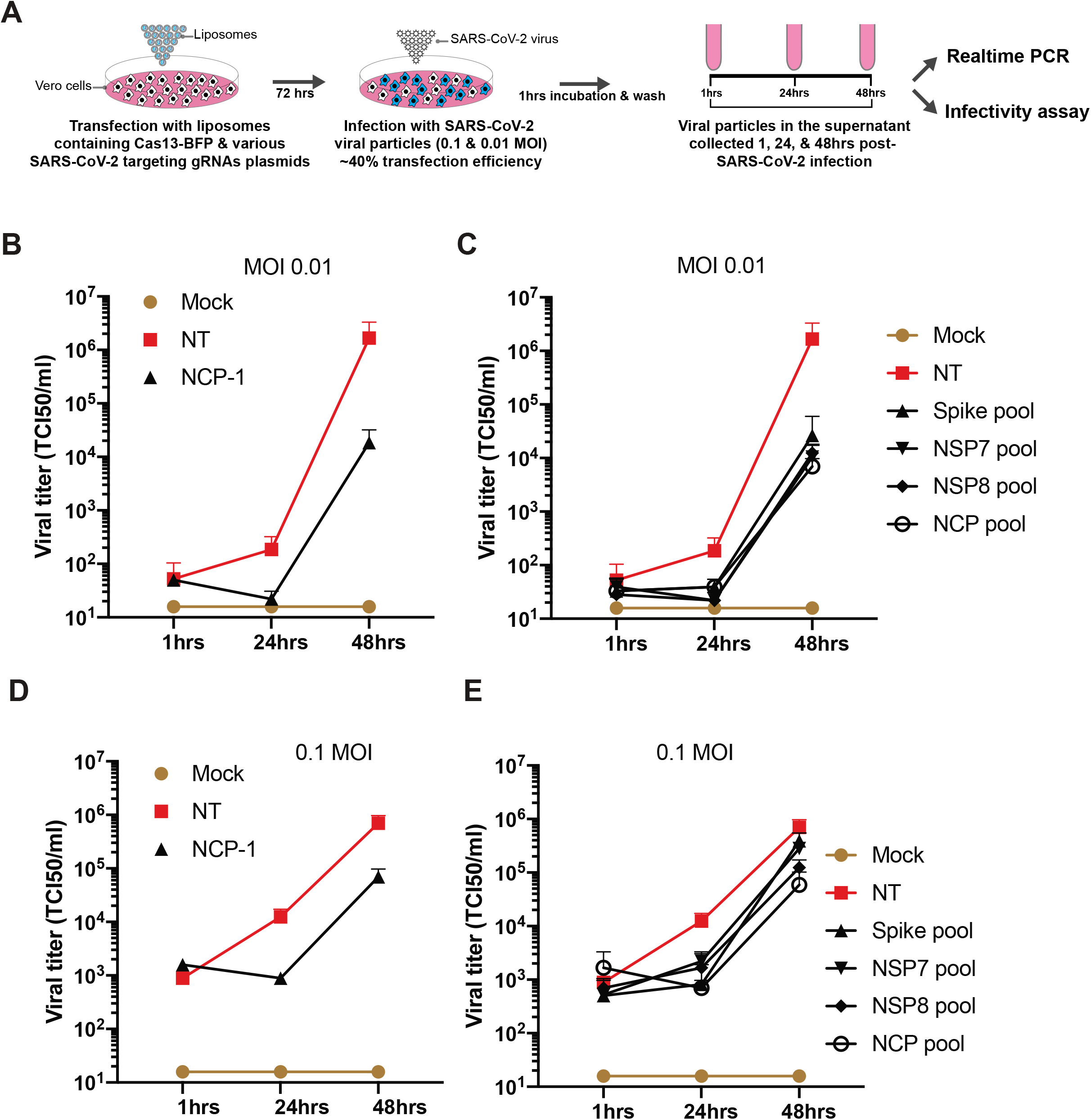
Silencing of replication-competent SARS-CoV-2 virus in infected VERO cells. (**A**) Schematic of infection assay to assess pspCas13b-mediated suppression of SARS-CoV-2 replication in infected VERO cells. VERO cells were transfected with liposomes containing pspCas13b-BFP and various gRNA constructs. 72h post-transfection, cells were infected with SARS-CoV-2 for 1 hour and the kinetic of viral replication was measured by assessing the infectivity of virus shed into the culture medium at 1, 24, and 48hrs post-infection (**B, D**) infectivity assay to evaluate the kinetics of viral replication in VERO cells expressing either NT or NCP-targeting gRNA1 at 0.01 (B) or 0.1 MOI. (**C, E**) infectivity assays to monitor the kinetics of viral replication in VERO cells expressing either NT or various pools of gRNA targeting NCP, Spike, NSP7 (RdRP subunit 7), or NSP8 at 0.01 (**C**) or 0,1 (**E**) MOI. N=2.

Suppl. Table 1

(https://docs.google.com/spreadsheets/d/12Vx8S4SSK3RY5He3gH6-yDDKJvovCVppI6-E0lX9lg/edit#gid=603752573)

**Suppl. Table 2**

(https://docs.google.eom/spreadsheets/d/1v58E-7YTMU1JnARHPs3MY-WybtDRGQhzznUfU9SfNQ/edit#gid=1093504085)

**Suppl. Table 3**

(https://docs.google.com/spreadsheets/d/1Mpl476AAzg9Q07Si3lBfxaDLBoeR0bLaRlbfDNaQgGk/edit#gid=936490712)

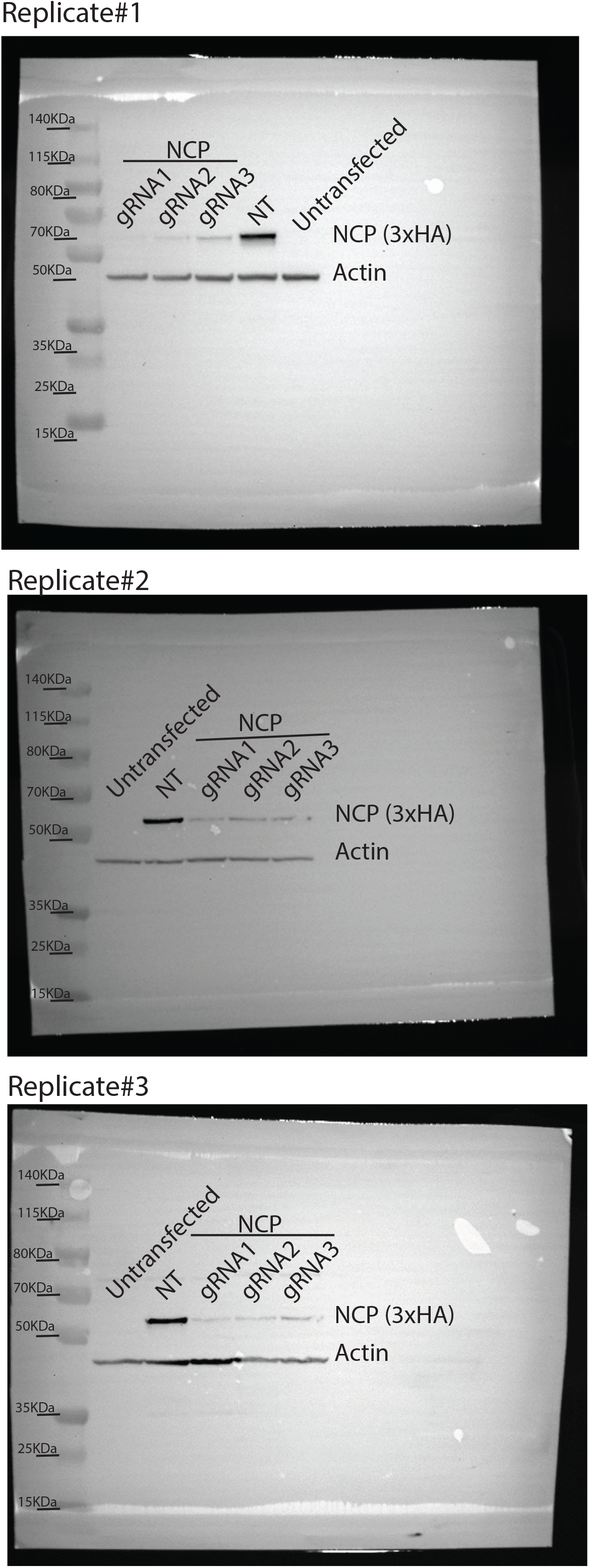

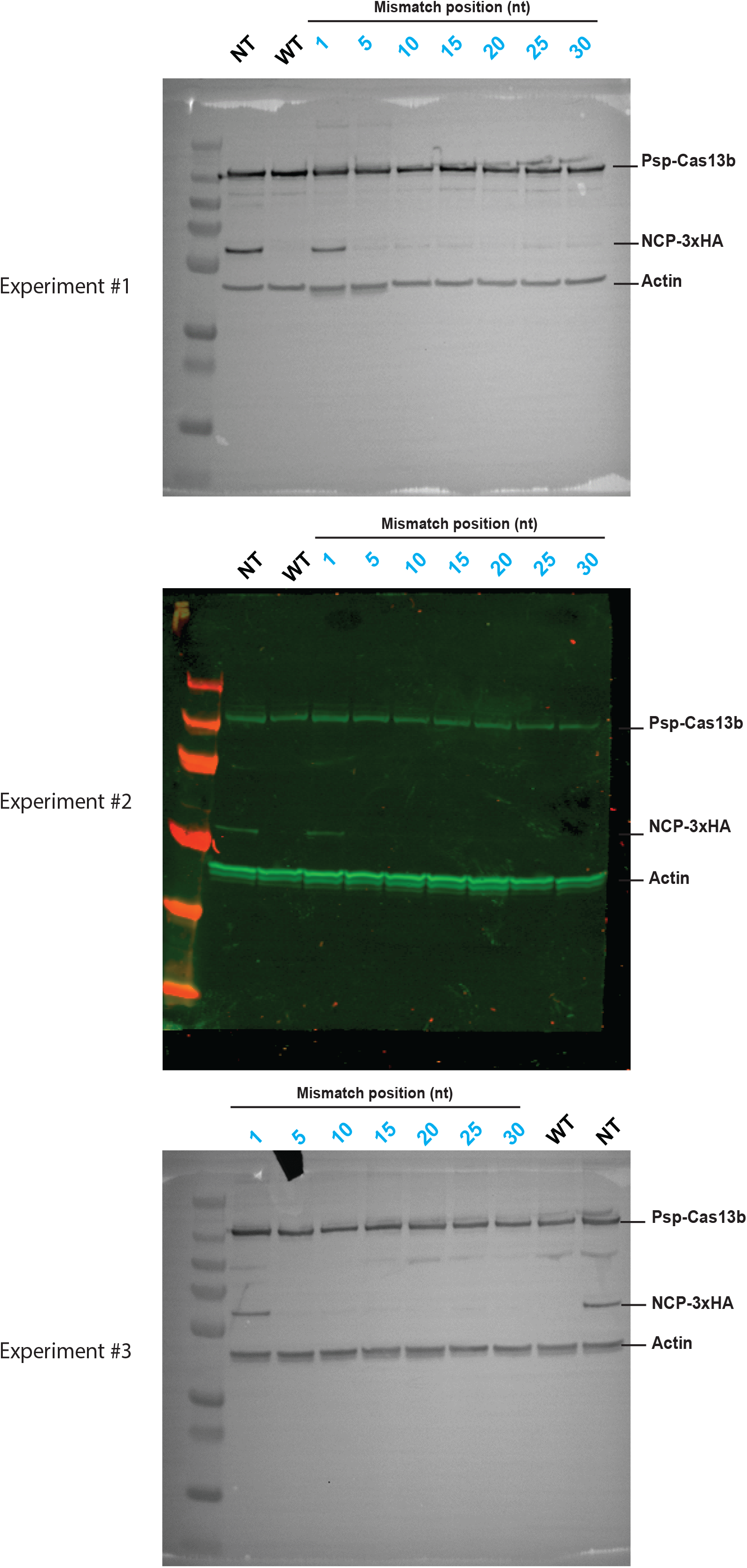

